# Defining the *in vivo* mechanism of air pollutant toxicity using murine stress response biomarkers

**DOI:** 10.1101/2022.10.05.510981

**Authors:** Francisco Inesta-Vaquera, Lisa Miyasita, Jonathan Grigg, Colin J. Henderson, C. Roland Wolf

**Author notes:** Corresponding author, James Arnot Drive, Jacqui Wood Cancer Centre. University of Dundee. Dundee, DD1 9SY. The authors declare they have no actual or potential competing financial interests.

## Abstract

**Background:** Air pollution can cause a wide range of serious human diseases. For the informed instigation of interventions which prevent these outcomes there is an urgent need to develop robust *in vivo* biomarkers which provide insights into mechanisms of toxicity and relate pollutants to specific adverse outcomes.

**Objective:** To exemplify the application of *in vivo* stress response reporters in establishing mechanisms of air pollution toxicity and the application of this knowledge in epidemiological studies and potentially in disease prevention.

**Methods:** Murine stress-reporter models (oxidative stress/inflammation, DNA damage and Ah receptor -AhR-activity) and primary mouse and primary human nasal cells were exposed to chemicals present in diesel exhaust emissions, particulate matter (PM) standards (PM_2.5_-SRM2975, PM_10_-SRM1648b) or fresh roadside PM_10_. Stress reporter activity was analysed by luminescence assays and histochemical approaches in a panel of murine tissues. Biochemical, genetic and pharmacological approaches were used to establish the mechanism of the stress responses observed. Pneumococcal adhesion was assessed in exposed primary human nasal epithelial cells (HPNEpC).

**Results:** Nitro-PAHs induced Hmox1 and CYP1a1 reporters in a time- and dose-dependent, cell- and tissue-specific manner. NRF2 pathway mediated this Hmox1-reporter induction. SRM1658b, but not SRM2975, was a potent inducer of NRF2-dependent Hmox1 reporter activity in lung macrophages. Combined use of HPNEpC and *in vivo* reporters demonstrated that London roadside PM_10_ particles induced pneumococcal infection in HPNEpC mediated by oxidative stress responses.

**Discussion:** The combined use of *in vivo* reporter models with HPNEpC provides a robust approach to define the relationship between air pollutant exposure and health risks. These models can be used to hazard ranking environmental pollutants by considering the complexity of mechanisms of toxicity. These data will facilitate the relationship between toxic potential and the level of pollutant exposure in populations to be established and potentially extremely valuable tools for intervention studies.

## Introduction

Air pollution in community, household and occupational settings is a major cause of non-communicable disease mortality and a leading contributor to global disease burden [1]. It comprises a complex mixture of gases and particulate matter (PM) whose composition is subject to geographic (spatial/temporal), atmospheric and anthropocentric (origin of source) variation [2-4]. PMs are classified according to their aerodynamic properties as PM_10_ : coarse particles less than 10 μm in diameter; PM_2.5_ : fine particles less than 2.5 μm; and PM_0.1_ (or ultrafine particles, UFP): smaller than 100nm [5]. The World Health Organisation (WHO) has issued maximum guideline values for exposures to PM_2.5_ as low as 5 μg/m^3^ (long term/annual mean) and 15 μg/m^3^ (acute/24h mean). It is clearly demonstrated that high PM exposure exacerbates a wide range of human diseases, including cardiovascular, cerebrovascular and respiratory disease (e.g. chronic obstructive pulmonary disease COPD), as well as Type 2 diabetes, lung cancer, pneumonia, chronic kidney disease, hypertension and dementia [6]. These studies have also increased our understanding of the complex biological effects of PM, including acute phase responses [7], hypertension [8], increased white blood cell and platelet counts [9], amongst others [10]. Despite an overall decrease of atmospheric pollution in developed countries, the prevalence of associated disease remains high. This is partly explained as the underlying biochemical and cellular processes involved are affected even at the lowest levels of exposure [11, 12].

There are major challenges associated to the study of environmental pollutants toxic properties, including the identification in complex mixtures of the constituents driving the observed health effects, the determination of toxic mechanisms involved and the extrapolation of laboratory data to interventions for vulnerable populations [13, 14]. For example, diesel exhaust particles (DEPs; a component of PM) are composed of large number of components, including carbonaceous aggregates with various proportions of metals and organic species adsorbed on their surface such as the polycyclic aromatic hydrocarbons (PAHs), nitro-derivatives of PAHs, oxygenated PAH derivatives (ketones, quinones, diones), heterocyclic organic compounds, aldehydes, and aliphatic hydrocarbons [15]. Mechanistically, the use of *in vitro* approaches has indicated that the deleterious health effects of DEPs are the consequence of an interplay between generation of oxidative stress and inflammation [16-19]. Protective antioxidant responses would predominate at low exposure doses, while inflammation and injury would occur at higher concentrations when cellular defences are overwhelmed [19, 20]. Using *in vivo* models, it has also been detected cytotoxic injury followed by an intense inflammatory process in the lung of mice models exposed to DEPs [21, 22]. However, there is a major need to develop novel tools that can accelerate the extrapolation of this data into the design of epidemiological studies to inform regulatory decisions. Part of this slow translation of findings is due to a lack of harmonisation between *in vivo* studies and their reliance on secondary markers of cell damage that are prone to artefacts, due to well-understood methodological problems. Current assays provide only a summary measure of damage averaged over the whole body at a particular timepoint and lack the cellular and tissue resolution required to draw meaningful interpretations of the responses observed.

To overcome these knowledge and methodological gaps, our lab has validated a panel of reporter models to detect *in vivo* the activation of major stress pathways linked to cellular toxicity, including oxidative stress/inflammation (HO1 triple transgenic reporter, HOTT), DNA damage (p21 reporter) or the AhR receptor activation (CYP1a1 reporter). These reporters provide easily measurable readouts of *in vivo* cellular responses to toxic insults, with high-resolution, and in a tissue- and cell-specific manner [23-25]. In these reporters, a short viral DNA sequence, known as a T2A sequence, is exploited to provide multiple reporter molecules to be expressed (separately) off the endogenous gene promoter. Recently, we have demonstrated the utility of these models as biomarkers for inorganic arsenic toxicity, a widespread water contaminant classified as a class 1 carcinogen [26]. We hypothesized that our stress reporters could be used as a complementary assay in environmental epidemiological studies to gain *in vivo* mechanistic insights into the associations observed.

## Materials and Methods

### Marylebone Road Particulate Matter

Marylebone Road (London, UK) PM_10_ (MR-PM_10_) was collected using a high-volume cyclone [1]. The cyclone was placed kerbside and within 1 metre of traffic, on various days throughout summer 2019 for 5 – 8 hours per day. Collections were pooled and stored at room temperature in a sterile glass container. Aliquots of MR-PM_10_ were diluted in Dulbecco’s phosphate-buffered saline (DPBS) to a final concentration of 1 mg/mL and stored as master stock at -20°C.

### Animals

All animals used in this study were supplied from the Medical School Resource Unit, University of Dundee, on a C57BL/6N background. Animals were subjected to the following husbandry conditions: mice were housed in temperature-controlled rooms at 21°C, with 45-65% relative humidity and 12h/12h light/dark cycle. Mice had *ad libitum* access to food (RM1 for stock mice; RM3 for mating females; Special Diet Services, 801010 and 801700 respectively) and water. Animals were regularly subjected to health and welfare monitoring as standard (twice daily). All cages had sawdust substrate and sizzle-nest material provided. Environmental enrichment was provided for all animals.

All animal work described was approved by the Welfare and Ethical treatment of Animals Committee of the University of Dundee. Those carrying out this work did so with Personal and Project Licenses granted by the UK Home Office under the Animals (Scientific Procedures) Act 1986, as amended by EU Directive 2010/63/EU. Animals in study plans were inspected regularly by staff trained and experienced in small animal husbandry, with 24-hour access to veterinary advice.

Animal numbers were guided by power calculations (G*Power; www.gpower.hhu.de), pilot experiments, and previous experience, and experimental design was undertaken in line the 3Rs principles of replacement, reduction, and refinement (www.nc3rs.org.uk.

### Animal studies design

Data in this paper was obtained using male mice, unless otherwise stated. Mice were 12-15wo littermates or age-matched to within 3 weeks of each other. All animals used in this study were heterozygous for the CYP1a1_KI_Cre, HOTT or p21 reporter allele unless otherwise specified. Animals were randomly assigned to control or treatment groups; analysts were not blinded to the identity of biological samples. At the end of studies, all animals were sacrificed according to Schedule 1 of the Animals (Scientific Procedures) Act 1986, by exposure to a rising concentration of CO_2_ and death confirmed by exsanguination, except animals for primary cells studies (cervical dislocation and confirmation of permanent cessation of the circulation).

### Mouse Treatments

3-nitrofluoranthene (Sigma; 405604) 1-nitropyrene (Sigma, N22959) and fluoranthene (Sigma; F807), Dexamethasone and Celecoxib were dissolved in corn oil with sonication for 1 h before administration by oral gavage (p.o.) at the indicated concentrations. Cisplatin (Sigma; C2210000) was dissolved in saline solution (0.9% NaCl in water) and administered by intraperitoneal injection (i.p.). N-acetyl cysteine (NAC) was dissolved in water. Particulate matter standards (SIGMA) was suspended in saline solution, sonicated and orophorayngeally deposited in anaesthetized mice. All animals (males) appeared healthy throughout the experiments and did not show clinical signs of illness or other potentially treatment-related symptoms. Organ weights were equivalent between control and treated animals at necropsy.

### Genotyping

Ear biopsies of mice 4–8 weeks old were incubated at 55°C for overnight in lysis buffer containing 75 mM NaCl (SIGMA, S9888), 25 mM EDTA (Millipore, 324503), 1% (w/v) SDS (SIGMA, S436143) and 100 μg ml−1 (39 U mg−1) proteinase K (Sigma; Cat. No. P6556). The concentration of NaCl in the reaction was raised to 0.6 M and a chloroform extraction was performed. Two volumes of isopropyl alcohol were added to the extracted supernatant to precipitate genomic DNA (gDNA). A 50 μl volume of TE buffer (10mMTris (SIGMA, 11814273001), 1mM EDTA, pH8.0) was added to the pellet, and subsequently gDNA was dissolved for 1h at room temperature and stored at 4°C until further use. The typical PCR sample consisted of a 25 μl volume containing 1.25U Taq DNA polymerase (Thermo-Scientific), 10mM dNTPs, Thermo Taq Buffer, 25mM MgCl_2_ and 10 pmol of the primers (HOTT reporter: HO1-KI Fwd, 5_-GCTGTATTACCTTTGGAGCAGG-3_; HO-1-KI Rvr, 5’-CCAAAGAGGTAGCTAATTCTATCAGG-3’); (p21 reporter: p21-KI Fwd, 5’-GCTACTTGTGCTGTTTGCACC-3’; p21-KI Rvr, 5’-TCAAGGCTTTAGGTTCAAGTACC-3’; Nrf2: Nrf2 Fwd, 5’-TGGACGGGACTATTGAAGGCTG-3’; Nrf2 As Rvr, 5’-GCCGCCTTTTCAGTAGATGGAGG-3’; Nrf2 LacZ Rvr, 5’-GCGGATTGACCGTAATGGGATAGG-3’); Tomato F – 5’-GGCATTAAAGCAGCGTATCC-3’); Tomato R – 5’-

### Tissue harvesting and processing for cryo-sectioning

Postmortem, tissues were rapidly harvested and processed by immersion fixation in 4% paraformaldehyde (brain, small intestine, skin) for 2h; 3% neutral buffered formalin (liver) for 3h; or Mirsky’s fixative (rest of tissues) for 24h. For cryosectioning, tissues were cryoprotected for 24h in 30% (w/v) sucrose in PBS at 4°C. Organs were embedded in Shandon M-1 Embedding Matrix in a dry ice-isopentane bath. Sectioning was performed on an OFT5000 cryostat (Bright Instrument Co.). With the exception of lung (14μm) and brain (20μm) sections, all sections were cut at 10μm thickness. Sections were kept at -20°C until further use.

### Imunohistochemistry

Tissue cryosections were allowed to dry (10min, RT) before rinsed in PBS (5 min., RT) and subsequently placed in a Coplan jar for antigen retrieval in citrate buffer (10mM sodium citrate pH 6, 0.05% Tween20; 9min, boiling temperature). Samples were washed in PBS, PBS-triton (0.03%) and PBS (each step for 5 min, RT). After washes, sections were blocked in 0.3% H2O2/MeOH (30min, RT), rinsed in PBS (5min, RT) and 2.5% anti-goat serum incubation (20min, RT). Cryosections were incubated with F4/80 antibody (Biolegend; Cat. no. 123101) in a humidified chamber (1/500 dilution in 5%BSA, overnight). After three washes in PBS – 0.05% Tween20 (5 min. each, RT), sections were incubated with peroxidase-conjugated secondary antibody (ImmPRESS goat anti-rat IgG; Vector laboratories, cat. no. MP-7444 ; 30min. RT). Finally, the sections were washed three times in PBS-Tween as before and DAB reaction (Vector laboratories; cat No SK-4100) time was monitored under microscope following manufacturer instructions. Immunolabeled sections were washed in running water for 5min. and mounted using Prolong (Invitrogen, P36930).

### Histochemistry

For *in situ* ß-galactosidase (Lac-Z) staining, tissue cryosections were rehydrated in PBS at room temperature for 15 minutes before being incubated overnight at 37°C in X-gal staining solution: PBS (pH 7.4) containing 2 mM MgCl_2_, 0.01% (w/v) sodium deoxycholate, 0.02% (v/v) Igepal-CA630, 5 mM potassium ferricyanide, 5 mM potassium ferrocyanide and 1 mg/ml 5-bromo-4-chloro-3-indolyl β-D-galactopyranoside. On the following day, slides were washed in PBS, counterstained in Nuclear FastRed (Vector Laboratories) for 4 min, washed twice in distilled water for 2 minutes and dehydrated through 70% and 95% ethanol (4.5 and 1 minute respectively) before being incubated in Histoclear (VWR) for 3 minutes, air-dried and mounted in DPX mountant (Sigma). Liver hematoxylin – eosin staining has been described previously (Inesta-Vaquera 2021) with minor modifications. Lung sections were stained with periodic acid–Schiff, according to the manufacturer’s recommendations (Sigma).

### Immunoblots

Whole-tissue protein extracts were prepared from snap frozen organs. Briefly, 300μl of lysis buffer (25mM Tris-HCl pH 7.4, 150mM NaCl, 5mM EDTA, 1% Nonidet P40, 0.5% sodium deoxycholate, 0.1% SDS) supplemented with protease inhibitors and phosphatase inhibitors (Roche, 04693132001; SIGMA, P5726) were added per 100mg of tissue and homogenized using a Polytron PT2100 benchtop homogenizer rotor. After 30 min on ice, the resulting lysate was centrifuged for 15 min at 13200 rpm on an Eppendorf tabletop centrifuge at 4°C. Supernatant was recovered and protein concentrations were measured by Bradford assay kit (Biorad). Cell lysates were prepared in loading buffer followed by sonication. SDS-PAGE and immunoblotting was carried out as previously described [27]. Antibodies used included hmox1 (Abcam, ab13243), b-gal (Promega, Z3781), p21 (BD Pharmingen, 556431), NQO1 (Ab2346), GAPDH (Cell signaling, 2118).

### Relative quantification of mRNA

Total RNA was isolated from 100mg of snap frozen tissues using Trizol Reagent (Invitrogen) followed by RNeasy kit (Qiagen), according to manufacturer’s instructions. cDNA was synthesized from 500ng of total RNA using the Omniscript RT-Kit (Qiagen), according to the manufacturer’s instructions. The RT-PCR mixes were prepared by mixing 1.3 μl of cDNA with 1 μl of TaqMan probe set, 10 μl of Universal PCR Master Mix (PerkinElmer Applied Biosystems) and 7.7 μl of nuclease-free water. For the real-time PCR analysis, the following pre-designed TaqMan probe sets (Thermofisher) in solution were used: Mm00516005_m1 (Hmox1); Mm00434228_m1 (IL1); Mm00446190_m1 (IL6); Mm99999915_g1 (GAPDH); Mm02619580_g1 (actin); Mm00443258_m1 (TNF). Data was acquired in a QuantStudio™ 5 – 96-Well 0.2mL Block system. The relative gene expression levels in different samples were calculated using the Comparative CT Method. The expression of actin or GAPDH mRNA were used as internal expression controls.

### Liver microsomes preparation

Liver samples were homogenised in SET buffer (1:5 w/v; 0.25M sucrose, 5mM EDTA, 20mM Tris-HCl, pH 7.4) supplemented with protease inhibitors (Roche) using a Polytron homogenizer. After a first centrifugation at 20000rpm for 20 min at 4°C, supernatants were transferred to 1.5ml ultra-Eppendorfs and centrifuged for 90min at 37000rpm in a Sorvall ultracentrifuge at 4°C. Supernantants were saved as the cytosolic enriched fractions and pellets were resuspended in 100μl SET buffer as microsomal enriched fraction.

### Cell culture

MCF7_AREc32 cells were cultured as described before (Wang et al., 2006). Briefly, cells were maintained in growing media (DMEM + Glutamax, 10% FBS, pen/strep 5ml) supplemented with 0.8mg/ml G418 (Roche, G418-RO) for maintenance or removed for exposure to chemicals. 96-well plates at 5000 cells/well.

To generate heterozygous HOTT reporter mouse embryonic fibroblasts (HOTThet MEFs) crosses between homozygous HOTT male mice and C57wt females were set up. MEFs were isolated from littermate mouse embryos at day E12.5. The heads and internal organs were removed from the embryos. The remaining tissues were then cut into small pieces, and incubated in trypsin (5 min, 37oC). Resulting individual cells were then plated in 96well plates and cultured in Dulbecco modified Eagle medium (DMEM) containing 10% serum, 2 mM l-glutamine, 1% penicillin/streptomycin and used up to passages 3 to 5.

Bone marrow precursor cells were extracted from mouse femurs under sterile conditions. Bone marrows were flushed with PBS, cells washed with PBS and resuspended in 10 plates/mouse containing 8ml of BMDM media (DMEM, 10% FBS, heat inactivated, 1mM Pyruvate, 1x GlutaMAX, Pen/Strep and supplemented with 10ng/ml murine M-CSF (Peprotech, 315-02) for plating onto non-tissue culture treated petri dishes. After 4 days, 5ml of media was added. After 7 days adherent macrophages were scraped from the petri dishes with versene (Gibco, 15040) and then plated into tissue culture grade plastic 96 well dishes and left for 24 h to re-attach before exposure to chemical compounds.

Mouse primary hepatocytes were isolated from male HOTThet or NRF2-KO_HOTThet reporter mice. Liver were perfused with perfusion buffer (10 mM HEPES pH 7.65, 137 mM NaCl, 7 mM KCl, 0.7 mM Na_2_ HPO_4_, and 0.5mM EDTA), followed by digestion buffer (137 mM NaCl, 7 mM KCl, 0.7 mM Na_2_ HPO_4_, 10 mM HEPES 5.1 mM CaCl_2_, pH7.65, type I collagenase from *Clostridium histolyticum* type IV; Sigma), and homogenate filtrated through a 70 *μ*m cell strainer. After centrifugation (400rpm, 5min, RT) hepatocytes were suspended in M199 + Glutamax (Invitrogen, Cat. no. 41150-020), 1% Penicillin/Streptomycin (Invitrogen; 15140), 0.1% BSA in PBS, 10% FCS, 10nM Insulin, 200nM 3,3’,5-Triiodo-L-thyronine (T3) (Sigma, Cat. no. T2877T3), 500nM Dexamethazone (Merck, 265005) and were allowed to adhere to six-well (5 × 10^5^ cells) or 96-well plates (1 × 10^4^ cells) for 4-6 hours in a CO_2_ incubator at 37°C. Plates had been pre-coated with 12.5 *μ*g/cm^2^ rat-tail collagen type I (Gibco, ThermoFisher Scientific, Perth, UK). Subsequently, medium was replaced with treatment media (M199+Glutamax, 1% Pen/Strep, 100nM Dexamethazone).

Human primary nasal epithelial cells (HPNEpC) were purchased from PromoCell® (Heidelberg, Germany), and maintained in airway epithelial cell growth medium, with supplement kit, Primocin (InvivoGen, San Diego, USA), and 10% FBS. Passage number was less than 5.

### Cell based assays

HOTT or ARE_luc luciferase reporter activity was measured using the Luciferase Assay System (Promega, E1500), according to the manufacturer’s instructions and luminescence quantified using the Orion II Microplate Luminometer (Berthold Detection Systems).

The CellTiter 96 AQueous One Solution Cell Proliferation Assay (Promega, G3582) was used to determine cell viability, as described by the manufacturer. Briefly, 20ul of the reagent was incubated with 100ul of the cell culture media (96 wells) and after 4h the sample absorbance was read at 490nm.

The cell glutathione content was determined using the GGSH/GSSH-Glo assay (Promega; V6611) according to manufacturer instructions and luminescence quantified as described above.

### Platelet-activating factor receptor expression

Airway cells were seeded overnight (2 × 10^5^ cells per well) into adherent cell culture plates and cultured with MR-PM_10_ for 2 hours before washing and detaching cells with trypsin. Cells were stained with an anti-PAFR primary antibody (1:200; ab104162 Abcam, Cambridge, UK) for 1 hour, with shaking at room temperature. To control for nonspecific background staining, a PAFR isotype control (1:200; ab172730, Abcam, Cambridge, UK) was included in all assays. The epithelial marker E-cadherin was also included to gate HPNEpC and exclude cell debris (1:100; ab1416, Abcam, Cambridge, UK). Cells were subsequently washed and stained with secondary antibodies conjugated to either Alexa Fluor 488 (1:3000; ab150077, Abcam, Cambridge, UK) for detection of PAFR/isotype expression or allophycocyanin (1:1500; ab130786, Abcam, Cambridge, UK) for detection of E-cadherin. Analysis was carried out on the BD FACS Canto II machine using BD FACSDiva software (BD Biosciences, Oxford, UK). PAFR is expressed as median fluorescence intensity (MFI).

### Pneumococcal adhesion

The Streptococcus pneumoniae type 2 encapsulated strain D39 (NCTC 7466) was purchased from the National Collection of Type Cultures (Central Public Health Laboratory, London, UK), grown to mid-logarithmic phase (OD600 = 0.4 to 0.6) in brain–heart infusion broth (BHI) (Oxoid, Basingstoke, UK) and stored at −80°C. The pneumococcal adhesion assay was conducted as previously described [2]. Briefly, airway epithelial cells were seeded overnight (2 × 105 cells per well) into adherent cell culture plates and exposed to MR-PM_10_ for 2 hours. Cells were thoroughly washed to remove PM and then S. pneumoniae D39 added for a further 2 hours to allow adhesion to HPNEpC. Cells were subsequently washed to remove non-adherent bacteria, lysed, and plated for colony forming unit count (CFU/mL). In this assay, CFU count reflects both the number of pneumococci adherent to the surface of cells and the number of intracellular bacteria.

The PAFR receptor blocker CV3988 (half maximal inhibitory concentration [IC50] 0.28 μM was added to pneumococcal adhesion assays at a final concentration of 20 μM to confirm a role for PAFR to the pneumococcal adhesion assay [28, 29]. The role of MR-PM_10_ mediated oxidative stress was determined by adding N-acetyl cysteine (NAC; Sigma-Aldrich) to adhesion assays at a final concentration 5 mmol/L, 30 minutes before and during exposure of cells to LU-PM_10_. N-acetyl cysteine was removed by thoroughly washing cells prior to performing the pneumococcal adhesion assay.

### Statistics

Data are summarised as median (IQR) and analysed by either Kruskal-Wallis test with Dunn’s multiple comparisons test, or Mann Whitney test. Analysis were performed using Prism 8 (GraphPad Software Inc., La Jolla, CA, USA), and p <0.05 considered statistically significant.

### Clinical chemistry

Terminal bleeds were collected in heparinized blood collection tubes (Sarstedt). Measurement of plasma Alanine aminotransferase (ALT) activity was performed at the Mary Lyon Centre Clinical Pathology Laboratory (MRC Harwell Institute, UK) on a Beckman Coulter AU680 Clinical Chemistry Analyser (Beckman Coulter, High Wycombe, UK) using Alanine aminotransferase (ALT) reagent (Beckman Coulter reference: OSR6107).

## Results

### 1. *Ex vivo* profiling of oxidative stress responses to individual components of diesel exhaust particles (DEPs)

In order to identify candidate DEP compounds for *in vivo* studies we performed cell-based screens with individual components (incorporating the main chemical classes found in diesel exhaust) and evaluated their capacity to activate different *in vitro* stress reporter systems. Initially we used the MCF7_AREc32 reporter line, which contains a luciferase gene construct controlled by eight copies of the ARE enhancer element. This reporter is activated by the oxidative stress sensor NRF2 [30]. The pro-oxidant tBHQ, used as a positive control, increased reporter activity ∼8 fold. The nitro-PAH 1 nitro-pyrene (1-NP) and 3-nitro-fluoranthene (3-NFA) increased the luciferase activity approximately 5 and 6 fold respectively (Figure 1A). These increases were maximal after 18h incubation and remained stable for 24h (Figure 1B, 1-NP; and Figure 1C, 3-NFA). At the concentrations used most of the compounds were non cytotoxic (Figure S1A). However, the oxy-PAHs: 1,4-naphtoquinone (1,4-NQ) and 9,10-phenanthrenequinone (9,10-PQ) were acutely cytotoxic at concentrations >1 μM with no induced ARE_luc activity (Figure 1A and Figure S1).

**Figure 1.**
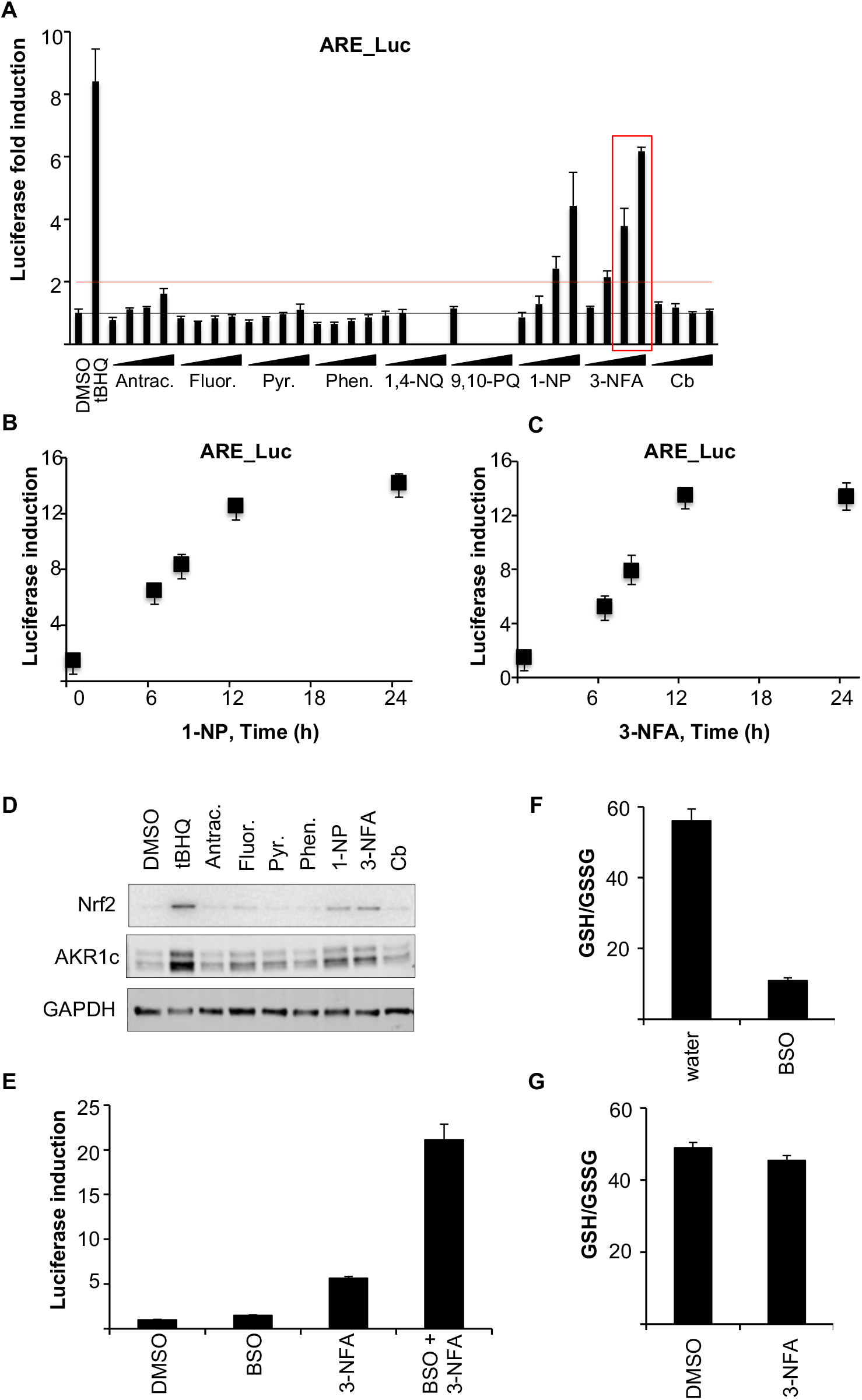
Identification of toxicity mechanisms of DEP compounds using human cell reporter assays. **A**. Luciferase activity was measured in MCF7-AREc32 cells exposed to increasing concentrations (1, 10, 25 or 50μM) of the indicated DEP compounds for 24h as described in the methods. **B and C** -as in **A**, but luciferase activity was measured at different time points after incubation with 50μM 1-NP (**B**) or 3-NFA (**C**). **D**. MCF7-AREc32 cells were exposed to indicate compounds (50μM) and 24h later whole cell protein extracts were immunoblotted with indicated antibodies. **E**. Cells were incubated with BSO (50μM), 3-NFA (50μM) or both for 24h and luciferase induction measured. **F**. MCF7-AREc32 cells were incubated with BSO (50μM) for 24h and GSH/GSSG ratio determined as indicated in methods. **G**. MCF7-AREc32 cells were incubated with 3-NFA (50μM) for 24h and GSH/GSSG ratio determined. Antrac.: anthracene; Fluo.: fluroanthracene; Pyr.: pyrene; Phen.: phenanthracene; 1,4-NQ.: 1,4-naphtoquinone; 9,10-PQ., 9,10-phenanthrenequinone; 1-NP., 1-nytropyrene; 3-NFA, 3-nitrofluoranthene; Cb, carbon black.

The nitro-PAHs (1-NP and 3-NFA), but not other tested PAHs, increased NRF2 protein levels (Figure 1D) and the expression of a bona-fide NRF2 target protein, AKR1C1/2. These results indicated that nitro-PAH compounds trigger a NRF2 response. To gain further insight into the mechanism of activation we modulated intracellular redox status in MCF7-AREc32 cells by pre-incubation with the glutathione depleting agent, buthionine sulphoximine (BSO) prior to exposure to 3-NFA. BSO alone did not increase reporter activity (Figure 1E) [30]. However, the combination of BSO with 3-NFA increased the induction from 4 to a 20-fold. Under these experimental conditions, BSO caused an 80% decrease in the GSH/GSSG ratio (Figure 1F). 50 μM 3-NFA alone for 24h only had a negligible effect in the GSH/GSSG ratio (Figure 1G).

We then established whether primary cell cultures derived from the HOTT reporter mice could be used to identify potentially toxic components of air pollution mixtures. In primary mouse embryonic fibroblasts (MEFs), luciferase activity was increased 3-fold when exposed to 20μM tBHQ for 24h (Figure 2A). When these cells were exposed to different DEP compounds, only 3-NFA (50 μM) caused a 2-fold induction of luciferase activity. Only oxy-PAHs were highly cytotoxic to MEF cells at concentrations >1μM (Figure S1B) and in the absence of reporter activity (Figure S1C).

**Figure 2.**
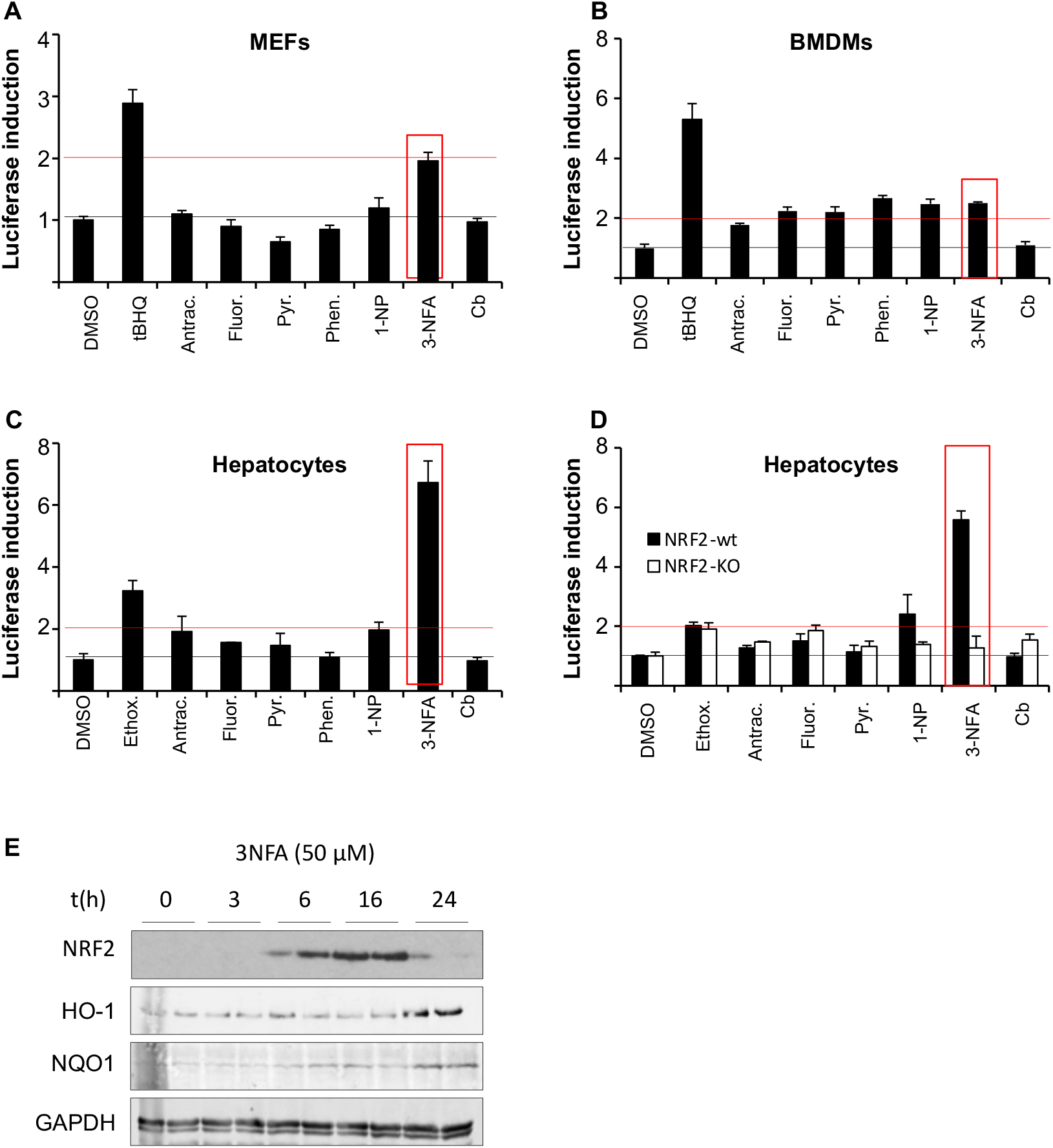
Identification of toxicity mechanisms of DEP compounds using primary cell reporter assays. **A**, mouse embryo fibroblasts (MEFS); **B**, bone marrow-derived macrophages (BMDM); and **C**, primary HOTT reporter hepatocytes were seeded in a 96-well plate and 12h later exposed to vehicle, 20μM tBHQ or 50μM of annotated DEPs and after 24h the luciferase activity was measured. **D**. -as in **C**, but cells derived from NRF2wt-HOTT reporter (black bars) or NRF2_KO-HOTT (white bars) reporter mice. **E**. Primary HOTT reporter hepatocytes were exposed to 3-NFA (50μM) for the indicated time points and whole cell protein extracts were prepared and immunoblotted for the indicated proteins.

We next studied the ability of DEP components to activate the HOTT reporter in immune cells. Primary bone marrow-derived macrophages (BMDMs) were exposed to DEP compounds and after 24h luciferase activity was measured (Figure 2B). The control compound tBHQ induced reporter activity by 5-fold and interestingly, there was a 2-fold increase in activity by a range of compounds, including both PAHs (fluoranthene, pyrene and phenanthrene) and nitro-PAHs (1-NP and 3-NFA) (Figure 2B). Carbon black did not induce the reporter expression in these cells.

We also tested the capacity of DEP compounds to activate the HOTT reporter in primary hepatocytes, a cell type with a high xenobiotic metabolising activity (Figure 2C). Cells exposed to 20 μM of the pro-oxidant increased luciferase activity on an average to 4-fold. When cells were exposed to 3-NFA (50 μM) a 7-fold increase in activity was measured. In contrast, exposure to 50 μM 1-NP only resulted in a 1.5 – 2-fold increase (Figure S1D).

In addition to Nrf2, Hmox1 is regulated by a number of other transcription factors[31]. To define the involvement of the NRF2 pathway in response to nitro-PAHs in primary hepatocytes we investigated reporter activation in hepatocytes from NRF2-KO_HOTThet mice (Figure 2E) [26]. In cells derived from NRF2-KO_HOTThet mice there was no increase of luciferase activity on exposure to nitro-PAHs. In addition, exposure of Nrf2 positive primary hepatocytes to 50 μM 3-NFA resulted in an increase in NRF2 protein at 6h and peaking at 16h. Also, the expression of NRF2 inducible proteins (HO-1 and NQO1) was also increased (Figure 2E). These data demonstrated a key role for Nrf2 in the responses observed.

### 2. Defining DEP toxic mechanisms *in vivo* using stress reporters

Based on the data obtained in our *in vitro* studies, we investigated the capacity of nitro-PAHs to activate *in vivo* stress responses [23, 24]. Mice carrying the HOTT reporter were treated with a single or 5 consecutive doses (at 24h intervals) of 3-NFA (50mg/kg). Twenty-four hours after the final dose tissues were harvested for histochemical and biochemical analysis (Figure 3 and Figure S2A). As previously described [23], there was a basal ß-galactosidase staining in the brain (hippocampus and cerebellum), kidney (tubular cells), and lung (bronchiole, respiratory epithelium). In mice treated with a single dose of 3-NFA a marked increase in LacZ staining was measured in liver (hepatocytes) and kidney (tubular cells) but not in any other tissue examined (Figure S2A). In mice exposed to repetitive 3-NFA doses, in addition to HOTT reporter expression in liver (hepatocytes and Kupffer cells) and kidney (tubular cells; Figure 3A), a striking increase in activity was observed in the heart (cardiomyocytes) and lungs (epithelial, muscle and bronchioles; Figure 3). There was no staining in any of the other tissues examined, including large intestine and brain (Figure S2A). Repetitive doses of 3-NFA did not induce significant changes in general toxicity markers, including plasma ALS/AST, H&E staining or mucopolysaccharides secretion in bronchioles (PAS staining; (Figure S3B, S3C and S3D respectively). NQO1 Western blot analysis confirmed an induction of NQO1 expression, but only in mice exposed to repetitive 3-NFA doses (Figure 3b).

**Figure 3.** Tissue specific susceptibility to 3-NFA exposure *in vivo* in HOTT reporter mice. **A**. HOTT^+/r^ male mice (n=3) were treated (p.o.) with vehicle or 50mg/kg 3-NFA once (single dose) or once daily for five consecutive days (repetitive doses). Tissues were harvested 24h later and *in situ β-*galactosidase assay performed in liver, heart, lung and kidney. **B**. Whole liver tissue extracts were prepared from mice in (**A**) and immunoblotted for indicated proteins. Black arrows indicate reporter activation.

To establish the role of NRF2 in the 3-NFA-induced activation of the HOTT reporter we treated reporter mice nulled at the Nrf2 locus with repetitive doses of 3-NFA (Figure 4A). Contrary to the robust 3-NFA induced reporter activity in the liver, kidney, lung, spleen and heart of NRF2wt-HOTT mice, reporter expression was largely abrogated in lung, heart and kidneys of Nrf2 null animals. Interestingly, modest reporter activation persisted in hepatocytes and Kupffer cells in these mice and Hmox1 reporter expression was activated in white pulp macrophages of the spleen by 3-NFA in both the NRF2 wt and null lines. To test the possibility that an inflammatory response could contribute to the reporter activation by prolonged 3-NFA exposure, we measured the expression of IL-6 and IL-1 genes in a 3-NFA responsive tissue (heart). 3-NFA did not increased the expression of these cytokines (Figure 4B), although there was an upregulation in the basal levels of these inflammatory markers (IL6 and IL1b) in NRF2-KO_HOTT reporter mice. This is in alignment with our previous observations (43) and reflects the interplay of NRF2 signalling and the immune responses.

**Figure 4.**
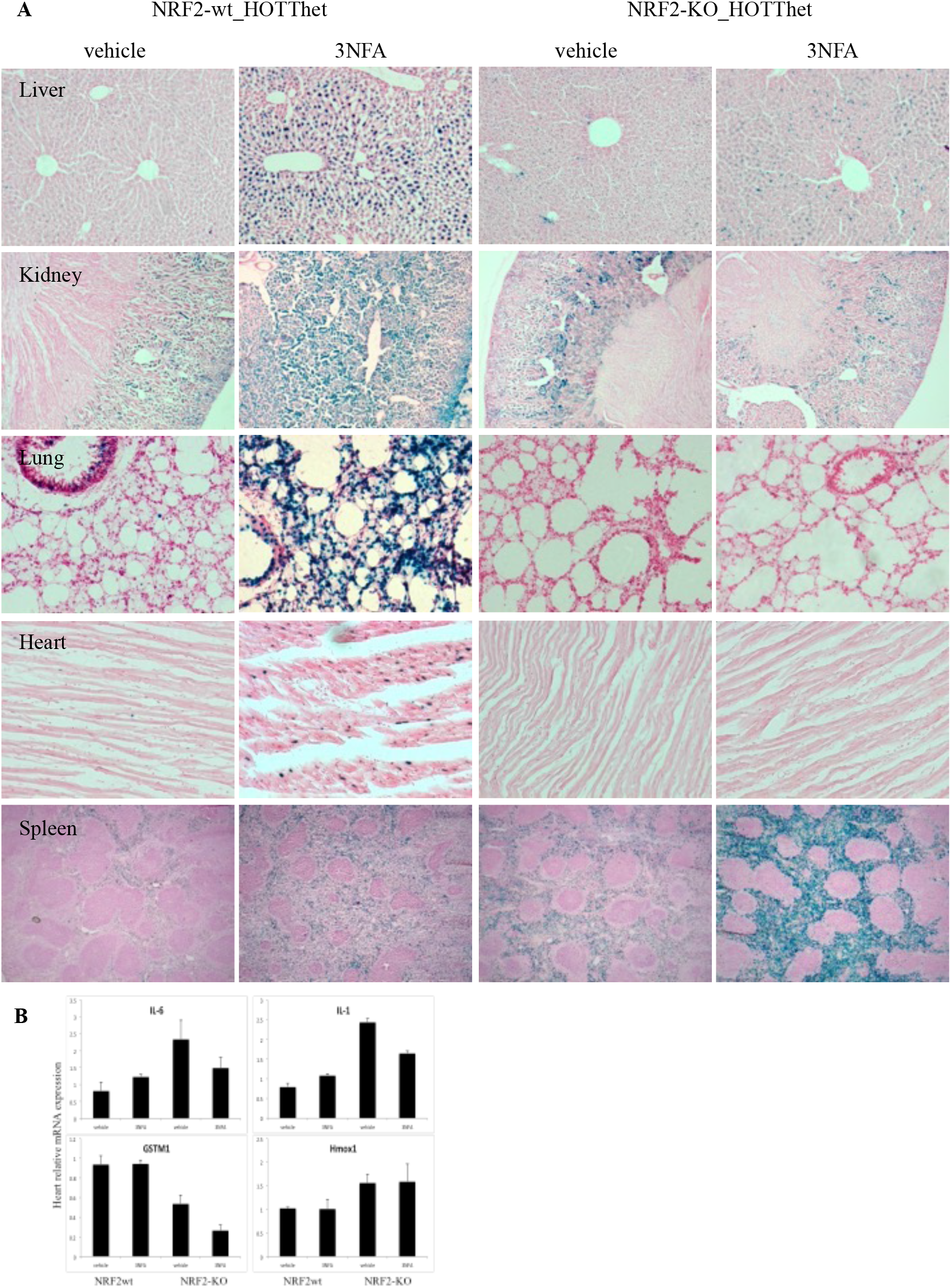
NRF2-dependent and independent HOTT reporter activation by 3-NFA *in vivo*. **A** -NRF2-wt_HOTT^+/r^ or NRF2-KO_HOTT^+/r^ male mice (n=3) were (p.o.) exposed to five repetitive doses of vehicle or 50mg/kg 3-NFA. Tissues were harvested 24h later and *in situ β*-galactosidase assay performed in liver, kidney, lung, heart and spleen. **B** relative mRNA expression of indicated genes in heart total RNA purified from animals in **A**.

We then extended these studies to another DEP compound, 1-NP. NRF2wt-HOTThet or NRF2-KO-HOTThet reporter mice were exposed to vehicle, single or repetitive doses of 1-NP at 50mg/kg (Figure 5A). Contrary to 3-NFA, a single dose of 1-NP did not activate the reporter in any of the tissues examined. Repetitive treatment only activated reporter expression in liver and sparsely in lung epithelial cells. No reporter expression by 1-NP was observed in NRF2-KO_HOTT reporter mice. Consistent with this finding, hepatic NQO1 protein was increased NRF2wt-HOTT mice after 5 consecutive doses but absent in NRF2-KO tissue (Figure 5B).

**Figure 5.**
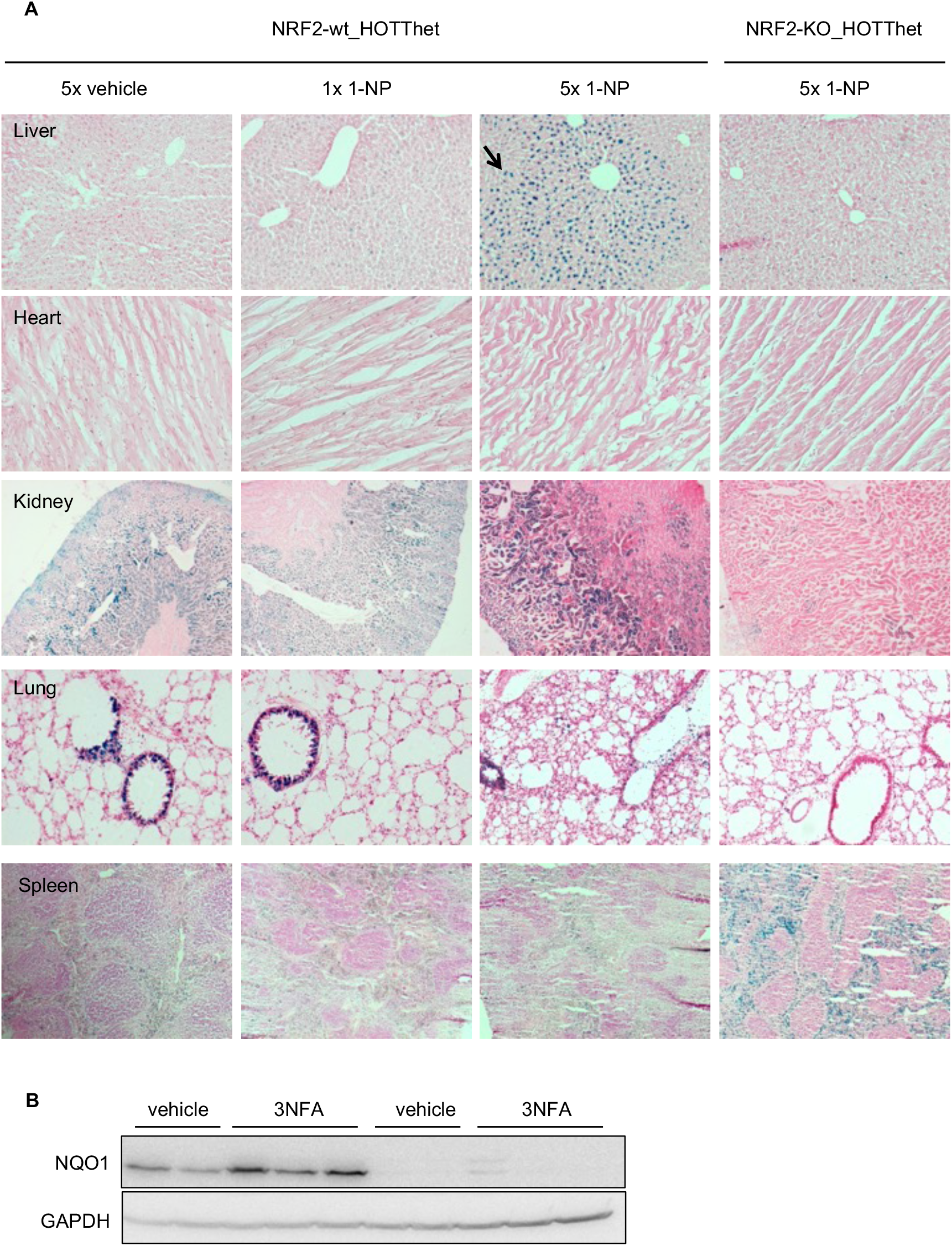
In vivo profiling of stress responses to 1-NP. **A**. NRF2-wt_HOTT^+/r^ or NRF2-KO_HOTT^+/r^ mice (n=3/group) were exposed (p.o.) to a single or five repetitive doses (once daily for 5 consecutive days) of vehicle or 50mg/kg 1-NP. Tissues were harvested 24h later and *in situ β*-galactosidase assay performed in indicated tissues. **B** liver whole tissue extracts were prepared and blotted against the indicated proteins. Black arrow indicates reporter activation.

Finally, we dosed HOTT reporter mice with fluoranthene following a repetitive doses protocol (Figure S3A). No activation of the HOTT reporter was observed in any of the tissues examined. AhR activation however was observed as hepatic CYP1a1 protein levels were increased (Figure S3B).

### 3. Studies on AhR and DNA damage-mediated responses

Nitro-PAHs can activate AhR as well as DNA damage responses [32-34]. We therefore examined the ability of 3-NFA to activate these pathways using a further set of *in vivo* reporters. AhR reporter mice (CYP1a1_KI_Cre-Tom^rep^) were treated with 5 daily doses of 50mg/kg 3-NFA. Basal activity was seen in some tissues, including liver, heart, kidney (Figure 6A), lung and spleen (not shown). On exposure to 3-NFA activation of the reporter was observed in hepatocytes only. In this experiment, AhR activation was confirmed by showing increased expression CYP1a1 and NQO-1 by western blot after the 3-NFA treatment (Figure 6C).

**Figure 6.**
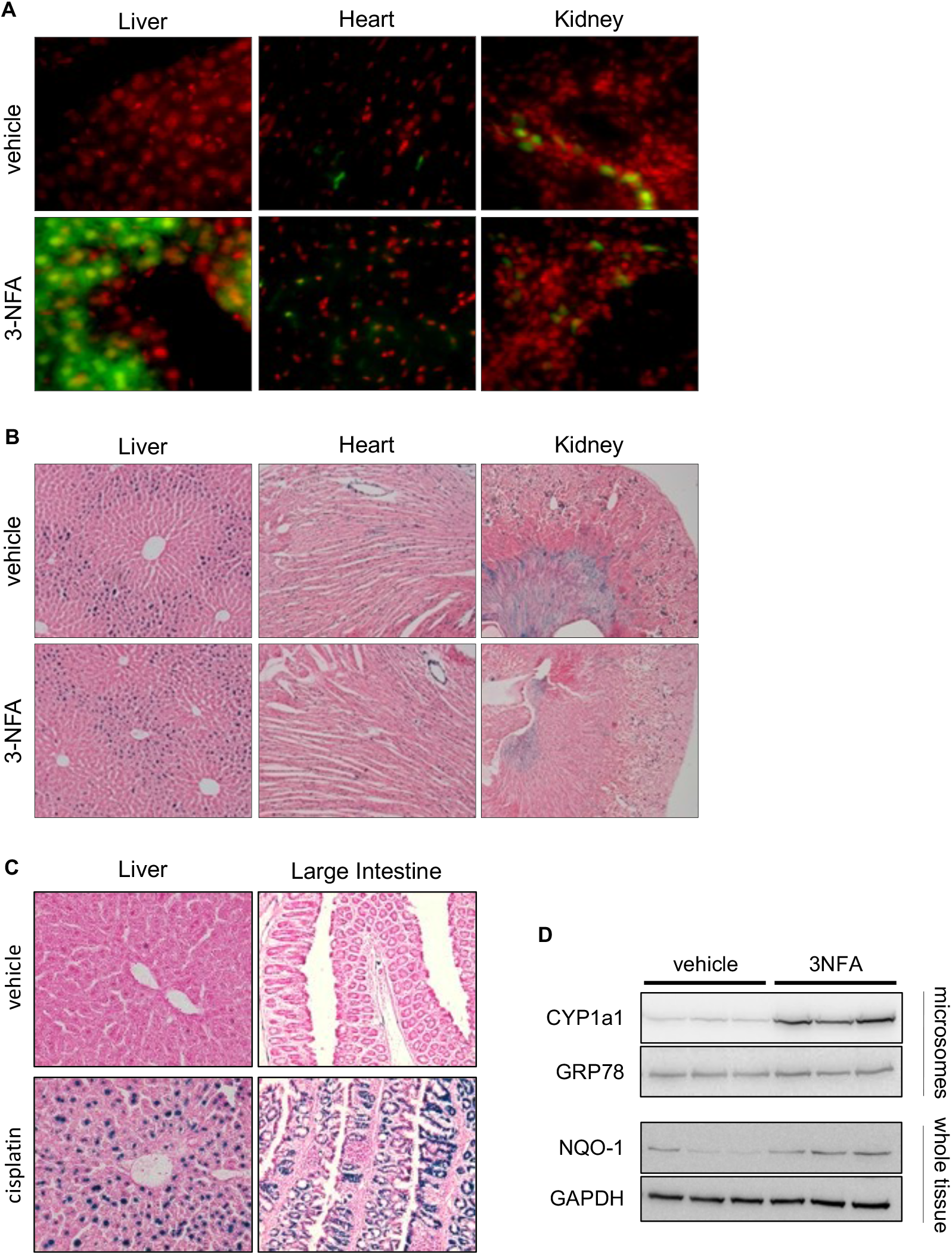
In vivo profiling of stress responses to 3-NFA using AhR and p21 reporter mice. **A**. CYP1a1_KI_Cre Tom^rep^ mice (n=3) were exposed (p.o.) to or five repetitive doses (once daily for 5 consecutive days) of corn oil (vehicle) or 50mg/kg 1-NP. Tissues were harvested 24h later and fluorescent microscopy used to detect mCherry signal (green) and DNA stain (Draq5, red). **B. p21** p21^+/r^ mice (n=3) were treated as in A. Tissues were harvested 24h later and *in situ β*-galactosidas*e* assay performed in indicated tissues. **C**. p21 mice were treated with saline solution (NaCl 0.9M) or cisplatin (10mg/kg, i.p.). Tissues were harvested 24h later and *in situ β*-galactosidas*e* assay performed in indicated tissues. **D**. Liver microsomes or whole tissue extracts derived from mice in **B** were blotted for indicated proteins.

We then investigated whether exposure to 3-NFA induced a DNA damage response *in vivo*. using the DNA damage-inducible p21 reporter [24]. Following 5 daily doses of 50mg/kg 3-NFA no changes over the basal expression of the reporter was observed in any of the tissues examined, including liver, kidney and heart (Figure 6B). Treatment with cisplatin, used as a positive control, resulted in marked p21 reporter activation in liver and large intestine (Figure 6D).

### 3. Pharmacological approaches to define toxic mechanisms involved in stress reporter activation by DEP components

To further define the mechanism of reporter activation by 3-NFA we employed a pharmacological approach. In the first instance, we tested the ability of the antioxidant NAC to modulate the reporter activation following 3-NFA treatment. We have previously shown that NAC at the concentration used (300mg/kg) does not increase the activation of the HOTT reporter [23, 26]. NAC administration completely prevented the induction of Lac-Z staining in the liver, but the reporter activity remained unaffected in the kidney.

In order to establish whether the Nrf2-independent reporter activity in the spleen induced by 3-NFA was due to an inflammatory response, NRF2-KO_HOTThet reporter mice were treated with either with 5 consecutive doses of 3-NFA alone or in combination with celecoxib (a selective Cox-2 inhibitor; Figure 7B) or dexamethasone (pan-NFKB signalling inhibitor; Figure 7C). Neither of these treatments attenuated reporter activation suggesting an alternative mechanism of induction. CYP1a1 induction in the liver (Figure 7D, E) and a reduction in plasma IL-1b were used as positive controls (Figure 7F).

**Figure 7.**
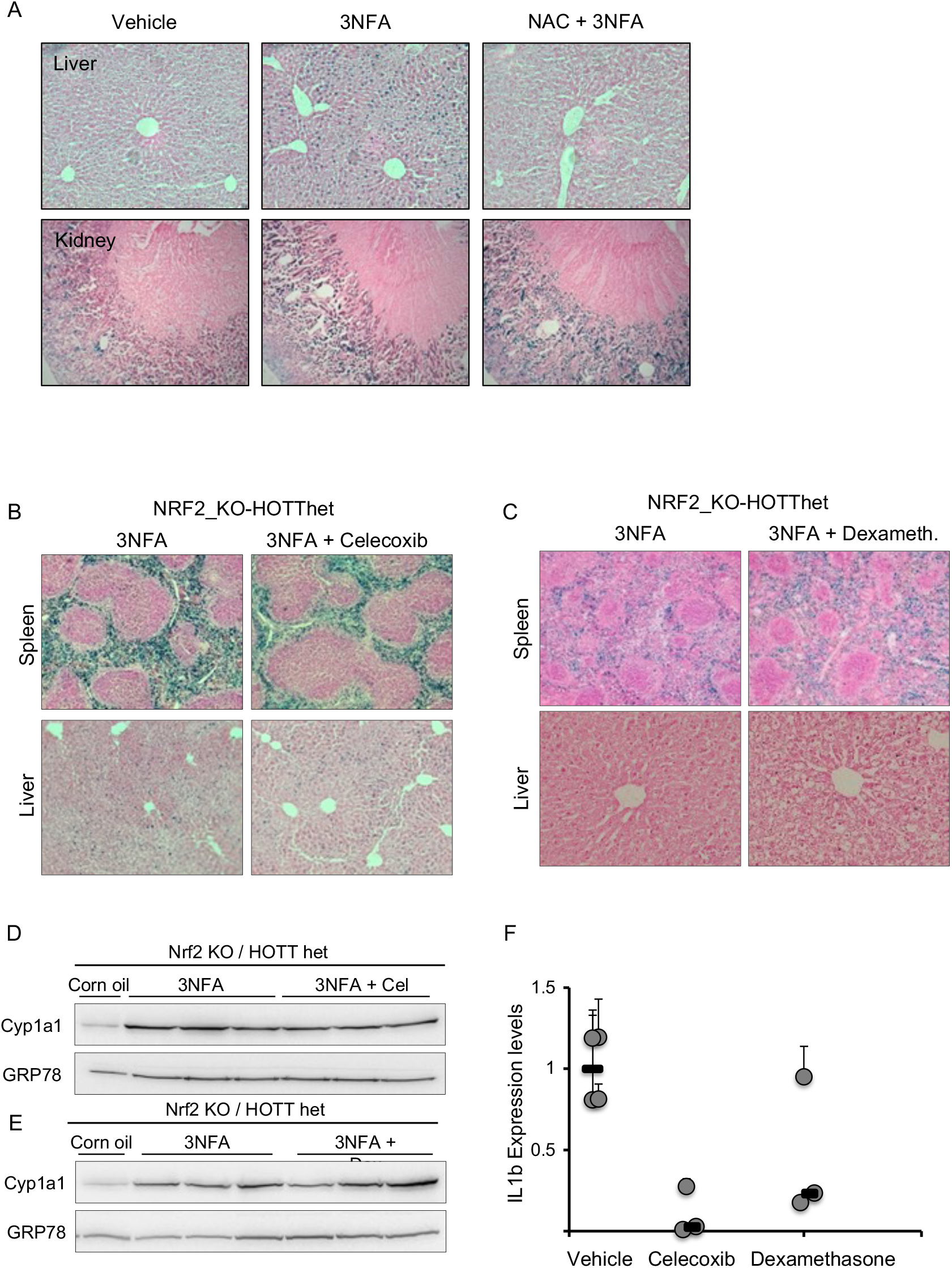
Mechanisms of HOTT reporter activation by 3-NFA. **A**. Triplicate HOTT reporter mice (n=3/group) were exposed to vehicles (corn oil p.o./ saline solution ip), 3-NFA (50mg/kg; po) or 3-NFA + NAC (300mg/kg, ip). Tissues were harvested 24h after dosing and *in situ β*-galactosidase assay performed in liver and kidney. **B, C**. Triplicate NRF2_KO-HOTT reporter mice (n=3/group) were exposed to 3-NFA (50mg/kg; 5 consecutive doses, once daily) and vehicle (corn oil) or celecoxib **(B)** (70mg/kg; 8 doses, once daily consecutive, starting 3 days before 3-NFA treatment) or in **(C)** dexamethasone (3mg/kg; 8 doses, once daily consecutive, starting 3 days before 3-NFA treatment) was used. 24h after treatments, tissues were harvested and *in situ β-galactosidase* assay performed in indicated tissues. **D, E**. Liver microsomes were prepared from animals in B and C, and immunoblotted for the indicated proteins. Corn oil sample (negative control) was extracted from animal in Figure 4. **F**. Total liver RNA was isolated and the relative expression of IL-1b was analysed by qRT-PCR.

### 4. Metabolic activation of air pollutants assessed in stress reporter mice

Metabolism at nitro-groups is an important mechanism of activation of substituted PAHs to toxic products [35]. Before apply *in vivo* stress reporters to investigate this mechanism of toxicity, we first exposed the metabolically competent MCF7_AREc32 cells to AhR agonists (TCDD 200ng/ml) 24h before and during exposure to increasing concentrations of selected PAHs and nitro-PAHs (Figure 8A). TCDD alone did not activate the luciferase activity, despite triggering a strong induction of CYP1a1 mRNA. 3-NFA also induced CYP1a1 in these cells, but to a lesser extent (Figure S4A). Treatment with a combination of TCDD and nitro-PAHs resulted in a marked additional 6-fold increase of luciferase activity compared to the activation by nitro-PAHs alone. TCDD did not increase reporter activity when cells were incubated with the PAHs fluoranthene or pyrene. In the light of these results, we investigated whether co-exposure of nitro-PAHs with other DEP components (i.e. PAHs) increased luciferase activity. Cells were treated with low doses of 1-NP or 3-NFA in combination with single PAHs for 24h. PAHs alone did not further induce the activation of the ARE_luc reporter by nitro-PAHs (Figure S4B).

**Figure 8.**
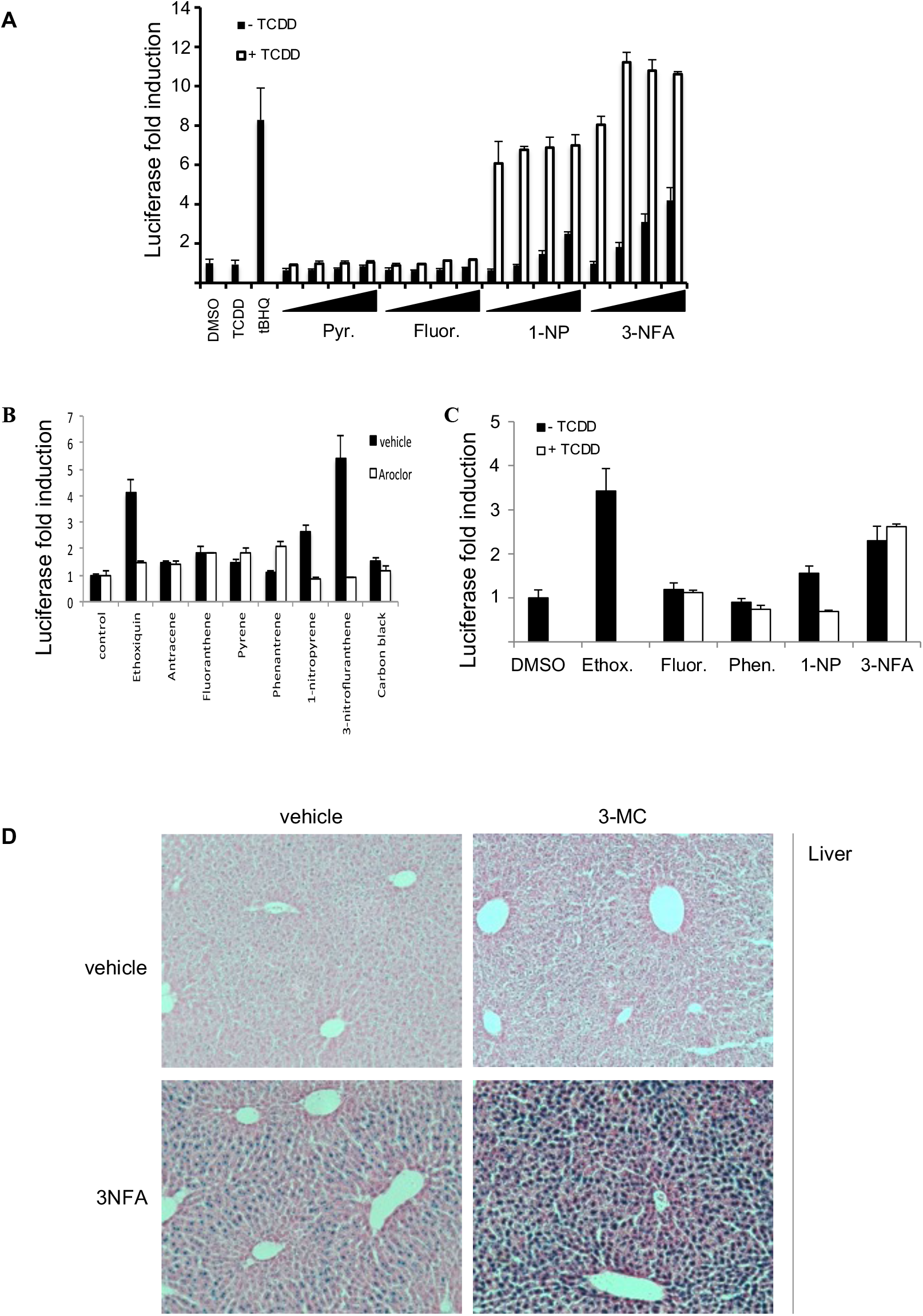
Contribution of metabolic activation to 3-NFA stress responses in vitro. **A**. MCF7_AREc32 cells were incubated with vehicle (black bars) or 100nM TCDD (white bars) for 24h before and during exposure to indicated DEP components (1, 10, 25 and 50 μM). **B**. Primary hepatocytes derived from HOTThet mice pre-treated with vehicle (black bars) or Aroclor-1254 (500mg/kg, ip; white bars) 4 days before cell isolation were incubated with indicated compounds (50 μM) for 24h before measuring luciferase activity. **C**. Primary hepatocytes derived from HOTThet mice were treated with vehicle (black bars) or TCDD (white bars) for 24h before incubation with indicated compounds (50μM) and 24h later luciferase activity measured. **D**. HOTT reporter mice (n=3) were treated with vehicle, 3MC (40mg/kg), 3-NFA (50mg/kg) or both. 24h later tissues were harvested and *in situ β-galactosidase* assay performed in liver

As a precursor to similar *in vivo* investigations, we carried out studies with primary hepatocytes derived from the HOTT reporter mice. Mice were administered vehicle or the AhR agonist Aroclor-1254, five days before cell isolation. Cells were then cultured in 96-well plates and exposed to a panel of DEP compounds for 24h (Figure 8B). AhR activation was confirmed by measuring Cyp1a1 protein (Figure S4C). Interestingly nitro-PAHs increased luciferase activity in hepatocytes derived from vehicle-treated reporter mice, however, on cells from Aroclor-1254 treated mice a reduction rather than an activation of the HOTT reporter with the compounds studies was observed. In addition, co-incubation of primary HOTTrep hepatocytes from untreated mice with TCDD and DEP compounds did not potentiate the activation of the luciferase reporter (Figure 8C). We confirmed the induction of Cyp1a1 expression by TCDD in whole cell lysates (Figure S4D).

The paradoxical data using primary hepatocytes can be explained by the complex balance between metabolic activation and detoxification as has been already described for certain PAHs [36]. We were therefore intrigued to establish the effects of the DEPs on reporter activity *in vivo*. Reporter mice were treated with the AhR agonist 3-methylcholanthrene (3-MC), (which does not induce a NRF2 response *in vivo)* and then to a single dose of 50mg/kg 3-NFA. Increased reporter expression was observed in liver and kidney of 3-MC-treated mice over untreated controls but not in any other tissues (Figures 8D, S4-E and S5).

### 5. Utility of reporter models to identify toxicity mechanisms of particulate mixtures and their correlation with responses in primary human nasal epithelial cells

In order to establish whether the use of *in vivo* reporters in mice identified toxic mechanisms of relevance to epidemiological studies in man, we dosed reporter mice with particulate matter representing different emission scenarios (Figure 9). Initially, HOTThet were exposed by oropharyngeal deposition to a single dose (250μg) or a repetitive dose (5x 50μg, once daily for 5 days) of the standard reference diesel particulate material (SRM 2975). Despite significant particle uptake in the alveolar epithelium, after 24h no reporter activation was seen in the lung or any other tissues examined (Figure 10A and not shown). We then tested a different pollutant mixture, SRM1648b (road dust PM_10_, Figure 9B) in NRF2wt-HOTThet and NRF2ko-HOTThet mice. Histological analysis showed that particles were mainly accumulated in the bronchioles, particularly in the terminal bronchiolar region and some extension up the bronchiolar tree (Figure 9B and C). Exposure to these particles induced reporter expression in a number of different cell populations in lung tissues, including bronchiolar cells and epithelial cells in both exposure protocols. Strikingly, these signals were significantly reduced in NRF2-KO mice, indicating that NRF2 plays a major role in the cellular responses associated with PM_10_ exposure (Figure 9C).

**Figure 9.**
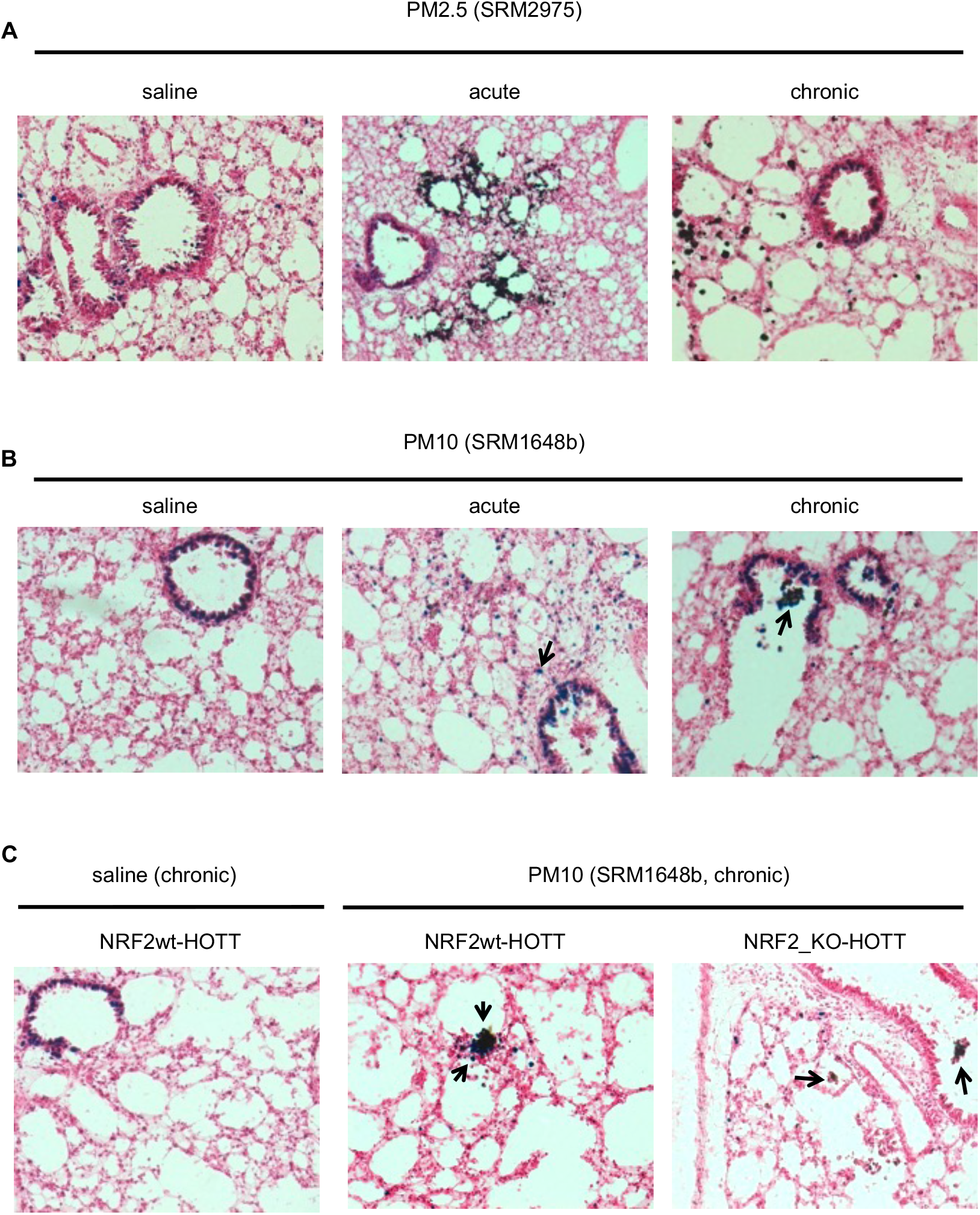
In vivo reporter activation by DEP particles. **A**. HOTT reporter mice (n=3) were instilled with saline solution (50μl) once daily for five consecutive days, a single dose of SRM2975 (50μl, 5μg/μl solution dispersed in saline solution), or five consecutive doses of SRM2975 (50μl, 1μg/μl, once daily for 5 consecutive days). 24h after treatments, tissues were harvested and LacZ staining was performed in lungs. **B**, as in (**A**), but PM_10_ SRM1648b was used. **C**, Triplicate reporter mice of indicated genotypes were instilled with five consecutive doses of saline or PM_10_ SRM1648b and tissues processed for LacZ staining as in (**A**). Black arrow indicates reporter activation.

**Figure 10.**
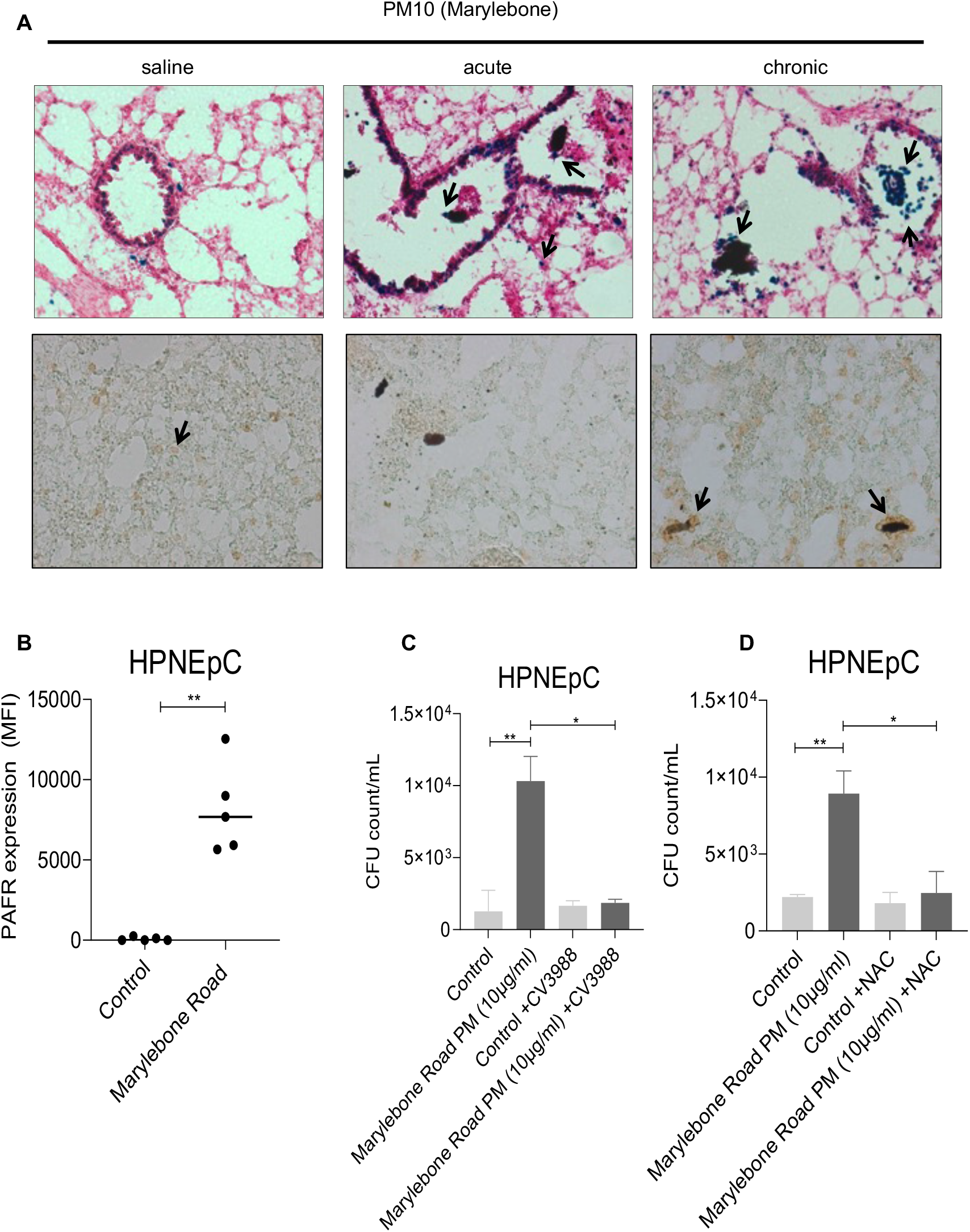
Utility of in vivo stress reporters to inform human studies. **A**. Triplicate HOTThet reporter mice were instilled with PM_10_ from Marylebone St. and LacZ staining was performed in lungs (upper panels). Cryosections derived from the same samples were subjected to F4/80 immunohistochemistry (lower panels). **B**. Effect of 10 μg/mL Marylebone Road PM10 (MR-PM10) on PAFR expression in human primary nasal epithelial cells. Data are from 5 separate experiments and summarised as median fluorescent intensity (MFI). Figure represents the median value and p values are calculated by Mann Whitney test. **C**. Effect of 10 μg/mL Marylebone Road PM10 (MR-PM_10_) on pneumococcal adhesion to human primary nasal epithelial cells (HPNEpC) and PAFR blocker CV3988 on MR-PM_10_ stimulated pneumococcal adhesion to HPNEpC. Data are from 6 separate experiments. Data are summarised as median IQR Kruskal– Wallis with *post hoc* multiple comparison testing. **D**. Effect of anti-oxidant N-acetylcysteine on MR-PM10 stimulated pneumococcal adhesion to HPNEpC. Data are from 7 separate experiments and are summarised as median IQR Kruskal–Wallis with post hoc multiple comparison testing. Black arrow indicates reporter activation.

Epidemiological studies have described an association of PM_10_ urban particulate matter exposure with increased vulnerability to bacterial pneumonia. We have previously shown that PM_10_ (**Leicester, Ghana**) has the capacity to increase pneumococcal adhesion as well as increase platelet-activating factor receptor (PAFR) expression in pulmonary cell lines. In order to establish whether the reporter mice could be used to evaluate toxicity mechanisms resulting from real time exposures, we collected London roadside PM (MR-PM_10_) using a high-volume cyclone. HOTThet reporter mice exposed to MR-PM_10_ revealed an increase in positive staining cells scattered through the lung parenchyma, particularly aggregated around inhaled particles (Figure 10A). The morphology of the cells was consistent with being alveolar macrophages and this was confirmed by an immunohistochemical analysis using the F4/80 macrophage marker. To define the human relevance of these findings we incubated primary human nasal epithelial cells (HPNEpC) with MR-PM_10_ (10 μg/mL). A significant increase in both PAFR expression (Figure 10B, p<0.01) and pneumococcal adhesion (Figures 10C-D, Yc, p<0.01) was observed. The PAFR receptor blocker CV3988 reduced MR-PM_10_ pneumococcal adhesion to baseline levels (Figure 10C, p<0.05). The antioxidant NAC inhibited MR-PM_10_ stimulated adhesion (Figure 10D, p<0.05) supporting a role for oxidative stress in this process. NAC had no effect pneumococcal adhesion in unstimulated cells.

## Discussion

We have used a panel of stress reporter mice to understand at a mechanistic level how environmental agents induce pathways associated with chemical toxicity. Although *in vitro* approaches can provide important insights into toxic mechanisms they have significant limitations in defining what actually happens *in vivo*. These include the physiological complexity of cell-cell interactions, metabolic activation, multicellular inflammatory responses (often considered to be a key factor in defining toxicity), crosstalk between tissues or *bona fide* replication of specific cell types. However, a major limitation of *in vivo* studies is the enormous amount of time, effort and cost involved in detailed pathological analysis of any cellular changes observed and also the fact that changes can only often be detected in the presence of overt toxic events. In order to circumvent this major challenge, we have for a number of years been developing a panel of reporter mice which reflect the activation of pathways directly associated with chemical toxicity. The activation of these pathways, including NRF2, p53 or AhR, provides insights into toxic mechanisms. The application of stress reporters to reflect toxic potential is widely accepted as reflected in the ToxCast 21 program or other environmental toxicology studies [37, 38].

In this paper we have exemplified the power of the *in vivo* stress reporters to understand toxic mechanisms of air pollution. Our studies provided a hazard ranking of toxic compounds and/or particles that can be used to inform epidemiology studies and identify health interventions aimed at reducing exposure to specific air pollutants. Additionally, through the use of a range of stress responsive reporters (e.g. oxidative stress, inflammation, AhR interactions and DNA damage), we obtained mechanistic insights into the toxicity of air pollutants. While the reporters used in this study represent some of the major mechanisms of toxicity related to the deleterious health effects of environmental exposure to air pollutants [39, 40], this approach can be extended to study additional mechanisms of toxicity such as those induced by endocrine disruptors.

Our approach involved a combination of *in vitro* and *in vivo* studies. We used primary cells derived from the reporter mice in *in vitro* screens to prioritize chemicals with a higher toxic potential for subsequent *in vivo* studies. We selected a panel of chemicals commonly identified in urban diesel exhaust particles, including PAHs, nitro-PAHs, oxy-PAHs and carbonaceous particles. The advantage of using primary cell assays derived from reporter mice compared to immortalised cell lines is that they do not bear genetic mutations in cell signalling that could compromise the interpretation of results. For example, the human lung cancer-derived alveolar cell line A549 often used in toxicology studies has constitutively high levels of antioxidant proteins as a result of the stabilization of NRF2 as a consequence of mutations in Keap1 [41].

Our *in vitro* studies demonstrated that different compounds found in diesel exhaust particles activate distinctive adaptive response pathways. Consistent with the literature, we found that PAH derivatives (i.e., oxy- and nitro-PAHs) have the highest capacity to induce oxidative stress, AhR responses and/or cytotoxicity. Interestingly, although both oxy- and nitro-PAHs have been linked to oxidative stress [42], the oxy-PAHs, 1,4 naphthoquinone, 9-10 phenanthrenequinone, did not activate an adaptive oxidative stress response in our studies. This could be ascribed to the differences in endpoints been measured. For example, 1,4 naphthoquinone increased the oxidation of DCFH-DA in melanoma cells at comparable concentrations used in our assays [43]. In contrast to quinones, we found that nitro-PAHs elicited a robust adaptive cell response in both human and murine cell reporter assays. Overall, our *in vitro* assays supported the conclusion that nitro-PAHs have a distinctive capacity to induce oxidative stress responses triggered by the NRF2 pathway. On this basis, we selected nitro-PAHs as chemical exemplars for our *in vivo* studies.

The application of multiple stress reporters for each individual compound or mixture *in vivo* allows a more complete assessment of toxic potential to be established. For example, we did not observe DNA damage responses where oxidative stress responses were observed. Furthermore, the use of multiple reporters allows cause and effect and relative dose and temporal responses to be established. We have shown for example that acute exposure to 3-NFA triggered NRF2-dependent Hmox1 reporter activation in hepatocytes and kidney tubular cells, indicative of an oxidative stress response. However, on chronic exposure activation was additionally observed in cardiomyocytes, lung (epithelial, bronchial and immune cells) and spleen (white pulp macrophages). We also demonstrated that compounds within a related chemical class (i.e. nitro-PAHs) can exhibit profound differences in the induction of toxic responses *in vivo*. For example, unlike 3-NFA, chronic exposure to 1-NP only resulted in a mild response in hepatocytes. These *in vivo* results highlight the importance of evaluating individual components in pollutants of concern to predict differential contributions to disease progression. They also identify potential caveats when assuming similar toxic properties of compounds based on physicochemical characteristics. For example, nitro-PAHs are grouped into a similar mode of action by which their bio-activation will induce oxidative DNA damage, formation of DNA adducts and mutagenesis. While this is indeed the case for 3-nitrobenzanthrone or 6-nitrochrysene [44, 45], (our *in vitro* and *in vivo* studies with 3-NFA do not support this. In fact, despite 3-NFA exhibiting a higher capacity to induce NRF2 responses than 1-NP, we did not observe a DNA damage response for either compound. These results are consistent with studies in laboratory animals, which demonstrated that 3-NFA is only an extremely weak carcinogen (<5% tumours after 100 weeks exposure to 1g of compound) [46]. Our findings provide evidence that understanding toxic potential at a mechanistic level is of key importance in the design of epidemiological studies. For example, in diesel mixtures 1-NP is found at higher concentrations than 3-NFA, 3-NFA metabolites are a much more relevant marker of toxic potential. Toxic potential is inevitably defined by the level of an individual compound, the duration of exposure and its potency as a chemical toxin.

A powerful application of *in vivo* reporter systems is to define through the use of genetic or pharmacological interventions the pathways affected by chemical exposure. For example, Hmox1 can be regulated *in vivo* by a number of different signalling pathways, including NRF2, heme and inflammation [23, 47]. In our study this was exemplified by the finding that induction of reporter activity was lost on a Nrf2 null background. Also pharmacological intervention, using the antioxidant NAC, attenuated our reporter activity induced by 3-NFA in the liver. Additional studies to define the mechanism of NRF2 activation by nitro-PAHs are in progress.

Metabolic transformation of PAHs, for example P450 mediated metabolic activation or redox cycling of quinones, can play a key role in defining the toxicity of chemicals and chemical mixtures. Currently, *in vitro* assays often used in epidemiological studies as a proxy for “toxic potential”, such as oxidative potential, do not take this key factor in defining chemical toxicity into account [48] [35, 45]. In the case of nitro-PAHs, two biochemical pathways are responsible for the metabolic activation. These involve cytochrome P450-mediated ring C-oxidation to epoxides, with subsequent rearrangement to nitro-pyrenols or hydration to dihydrodiols; and through nitro-reduction by reductases (e.g. P450 reductase, NQO1/AKR1C, and to a lesser extent P450s CYP1A1/2) to nitroso-PAH, N-hydroxy-amino-PAH or amino-PAH [49, 50]. In our studies we clearly demonstrated that AhR activation both *in vitro* and *in vivo* led to a marked increase in reporter activation after 3-NFA exposure. CYP1A1 expression is highly regulated by AhR, and indeed CYP1a1-mediated generation of electrophiles has been intimately linked with the activation of NRF2 by PAHs [51]. Our data also demonstrates that the metabolic activation and toxic potential of 3-NFA is dominant in relation to the potential AhR induction of antioxidant genes resulting in detoxification [52]. Which mechanism predominates will in all likelihood be both compound and dose dependent.

We further propose that the use of mechanistically informative *in vivo* reporters provide a more informed approach in epidemiological studies to define the relationship between air pollutant exposure and health risks. In our study, we have exemplified this by testing different reference particulates *in vivo* which allowed their relative toxic potential to be established. Interestingly, SRM1658b (PM_10_) activated the Hmox1 reporter in pulmonary macrophages in a NRF2-dependent manner. NRF2 activation in macrophages is associated with an anti-inflammatory function which limits the interferon response [53]. We then carried out a combination of *in vivo* reporter assays and PAFR expression and pneumococcal adhesion from exposed human primary nasal cells in subsequent studies of a freshly collected PM. For the first time, we collected London Marylebone Road roadside PM_10_ using a high volume cyclone [54], which has the advantage of collecting PM as dry respirable powder. Marylebone Road PM_10_ was found to also induce the Hmox1 reporter expression in alveolar macrophages, indicating an initial role for oxidative stress in the toxicity of these particles. Informed by the in vivo reporter assays, we used the antioxidant N-acetylcysteine to show that oxidative stress is responsible for the increased pneumococcal adhesion to primary nasal cells [55]. Therefore, these studies confirmed the capacity of our *in vivo* reporters as a predictive tool to study air pollutants of concern in exposed populations.

In summary, we illustrate the power of *in vivo* stress reporters to understand the mode of action of compounds and mixtures associated to environmental pollution. Our results exemplify how such experiments can be used to guide epidemiological studies. We envisage that the combination of *in vivo* reporter mice with human disease mouse models will be particularly useful to understand the extent to which environmental pollution exacerbates disease pathogenesis in vulnerable populations. In addition, the use of reporter mice crossed into mice humanised for pathways of drug metabolism e.g. P450s [56] may further increase the relevance of data obtained to man and allow PK/PD models of toxicity to be developed.

## Acknowledgements

This research was funded by a Medical Research Council Project Grant (MR/R009848/1) grant awarded to CRW/CJH/JG and a NERC SPF-grant (NE/W002213/1) awarded to CRW.

We thank members of the Medical School Resource Unit staff, Ngaire Dennison (Named Veterinary Surgeon) and University of Dundee Biological Services for technical support and ensuring animal welfare at all times. We also thank Tanya Frangova, Febe Ferro and Cheryl Wood for technical assistance.

## Figure Legends

**Figure S1.**
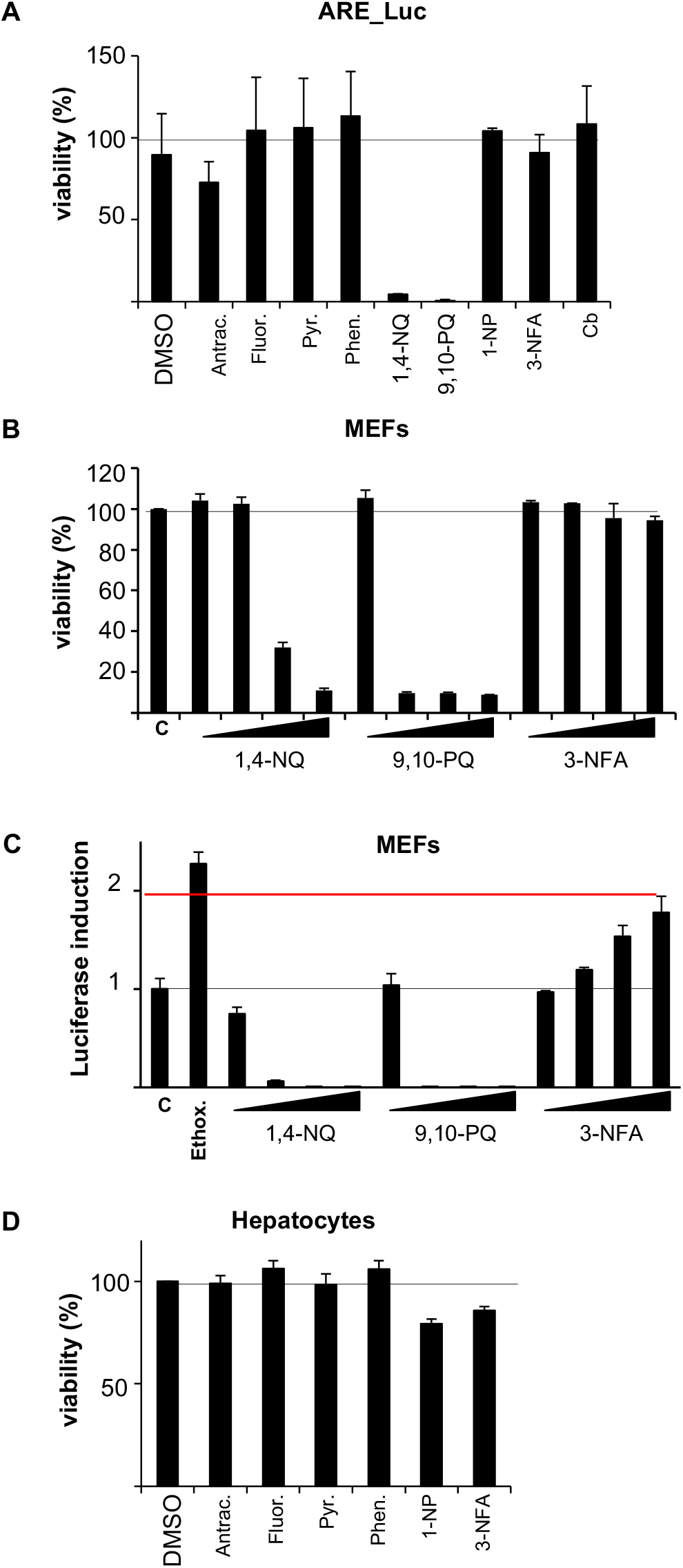
Cytotoxicity of DEP compounds. A. MCF7-AREc32 cells were exposed to 50μM of the indicated DEP compounds for 24h and cell viability measured. B. MEF cells were incubated with annotated compounds at 1, 10, 25 and 50μM for 24h and cell viability **(B)** or luciferase activity **(C)** was measured. **D**, as in **A**, but primary hepatocytes were analysed.

**Figure S2.**
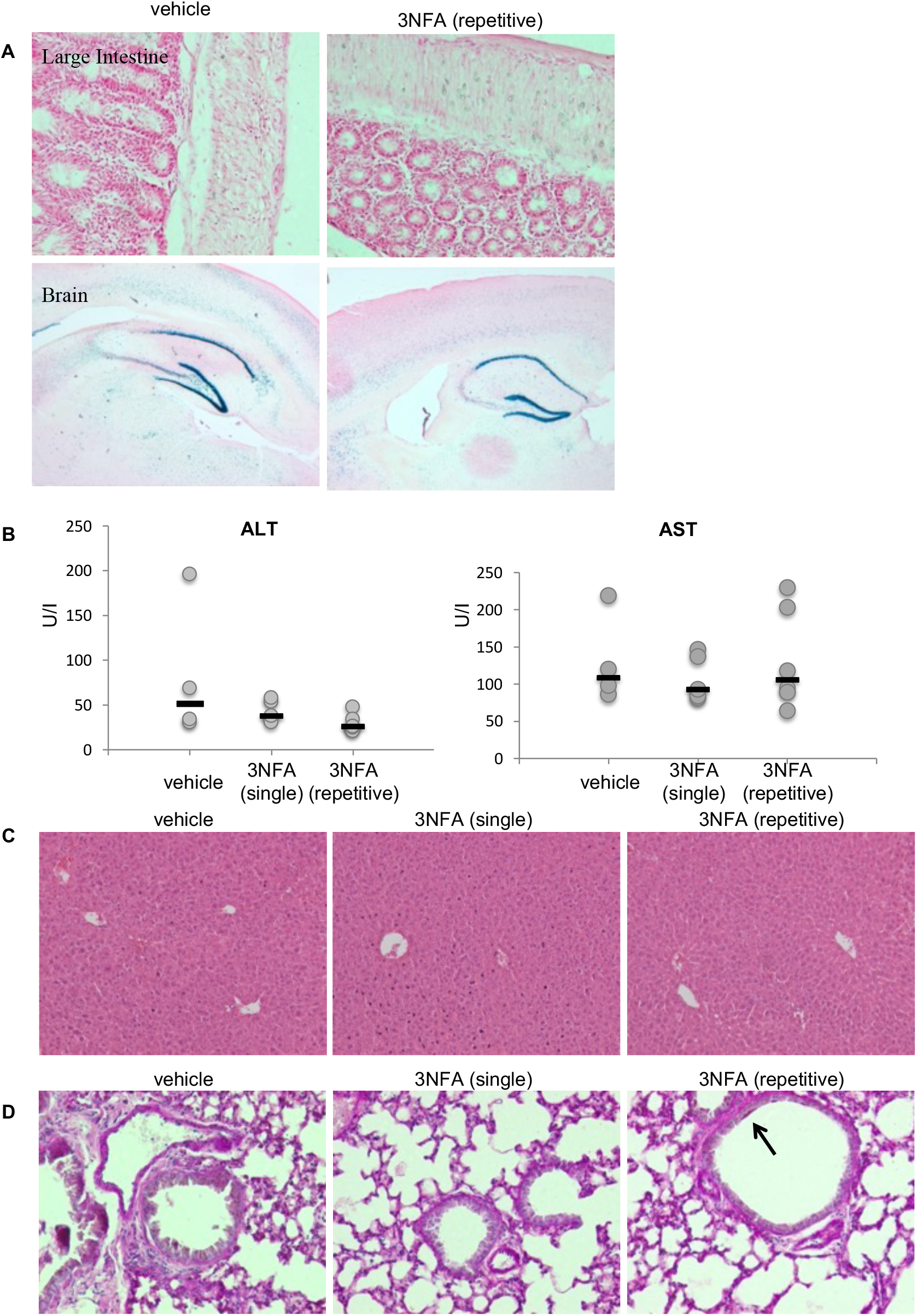
Tissue specific susceptibility to 3-NFA exposure *in vivo* in HOTT reporter mice. Samples were extracted from animals in Figure 3, but LacZ staining was performed in large intestine or brain (**A**), plasma levels of ALT and AST determined (**B**), liver sections stained for H&E histochemistry (**C**) and PAS staining performed in lungs (**D**). Black arrow indicates accumulation of mucopolysaccharides.

**Figure S3.**
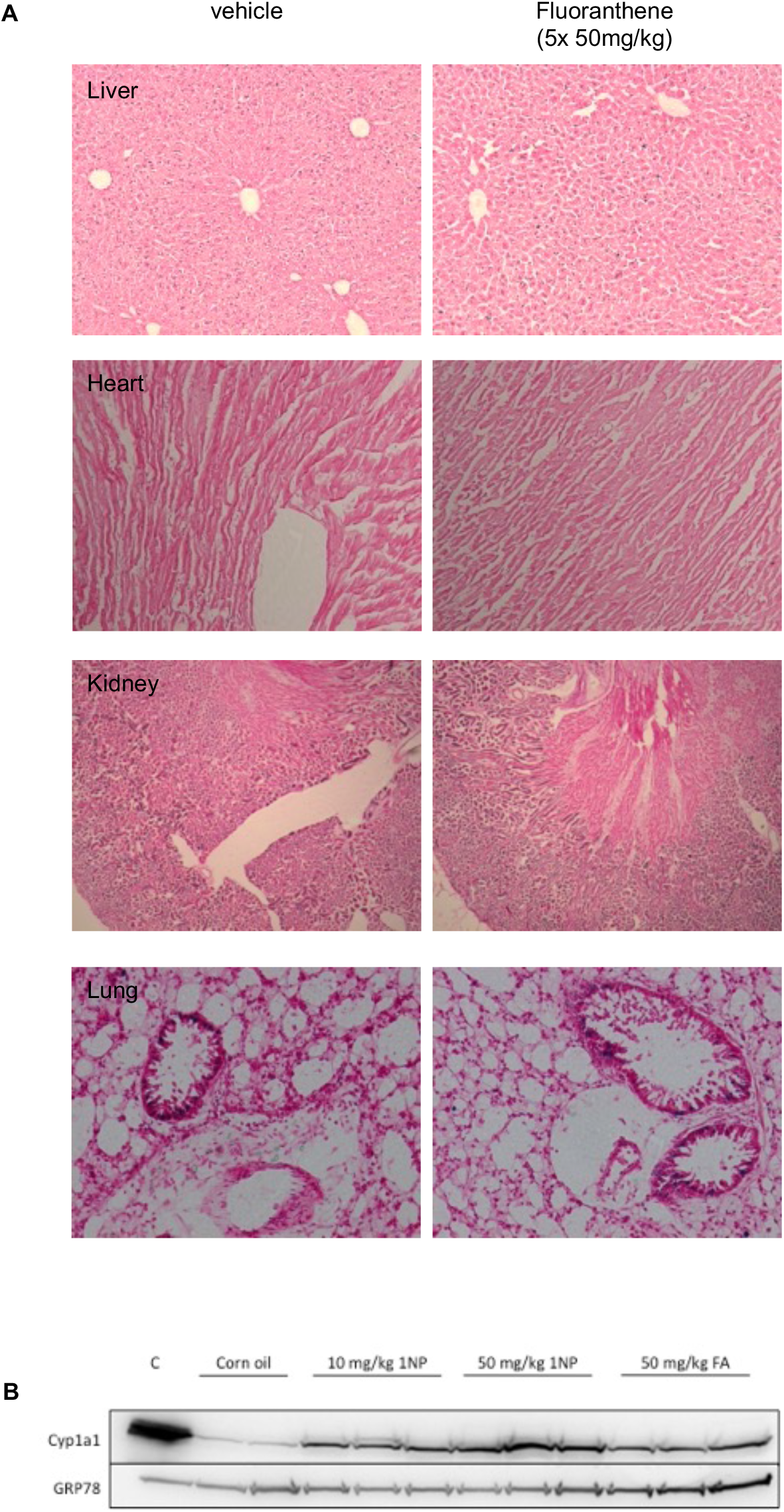
Fluoranthene do not activate the HOTTrep *in vivo*. **A**. Triplicate HOTT reporter mice were exposed to repetitive doses of vehicle (corn oil), or repetitive (50mg/kg, 5 consecutive doses, once daily). Tissues were harvested 24h after last dose and *in situ β*-galactosidase assay performed in indicated tissues. **B**. Liver microsomes derived from mice treated with indicated concentrations of 1NP or FA were immunoblotted against indicated proteins.

**Figure S4.**
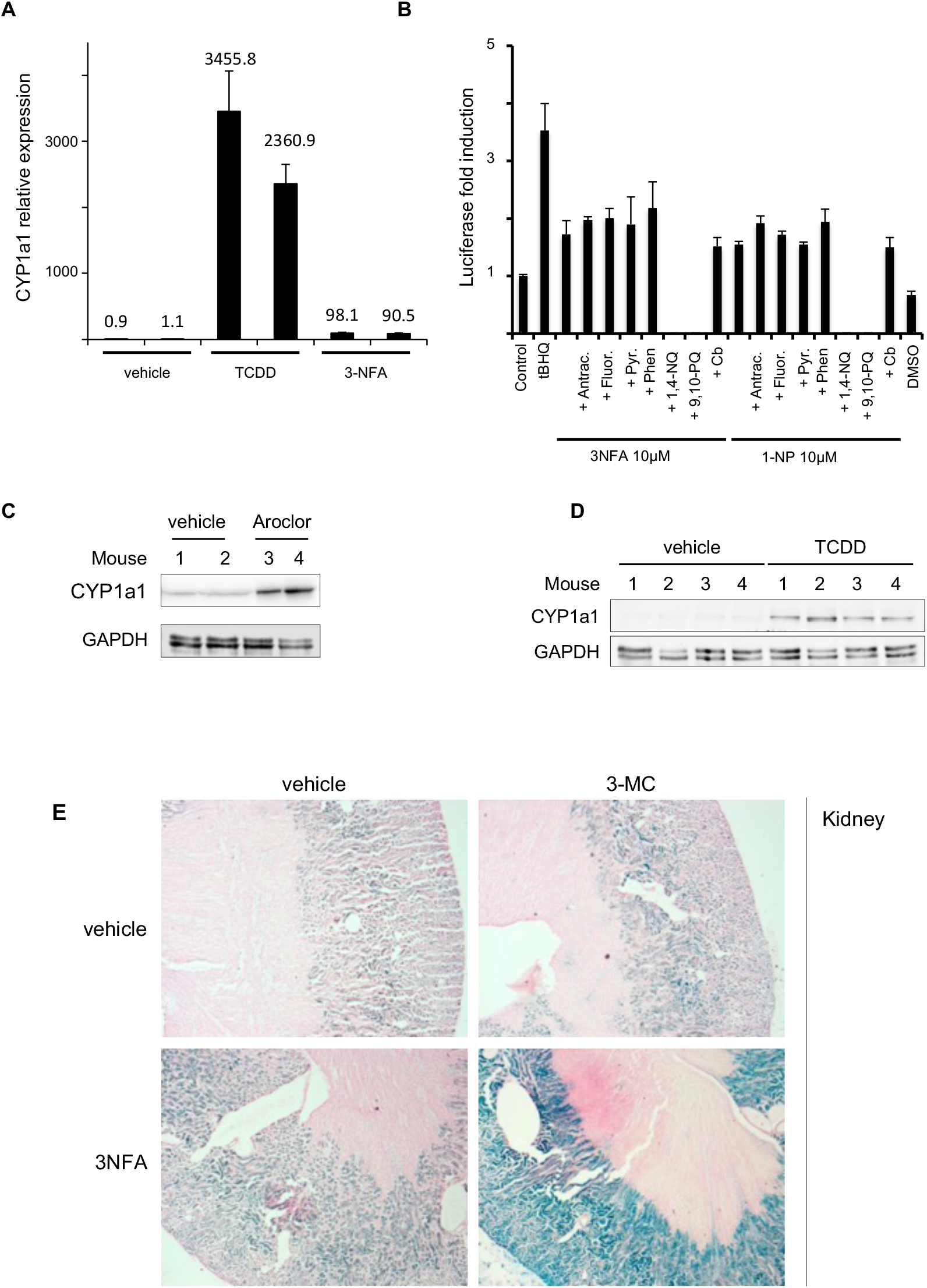
In vivo metabolic activation enhances oxidative stress responses to 3-NFA. **A**. MCF7_AREc32 incubated cells were incubated with vehicle, 100nM TCDD or 50μM 3-NFA for 24h before total RNA was isolated and the relative expression of CYP1a1 was analysis by qRT-PCR as described in methods. **B**. MCF7_AREc32 cells were treated with 10μM of indicated nitro-PAH alone or in combination with 10μM of indicated PAHs and 24h later luciferase activity was measured. **C**. Whole cell protein extracts were prepared from primary hepatocytes in Figure 8B and immunoblotted for CYP1a1 and GRP78 proteins. **D**. As in **(C)**, but primary hepatocytes used in Figure 8C. **E**. As in Figure 8D, but kidney sections are shown.

**Figure S5.**
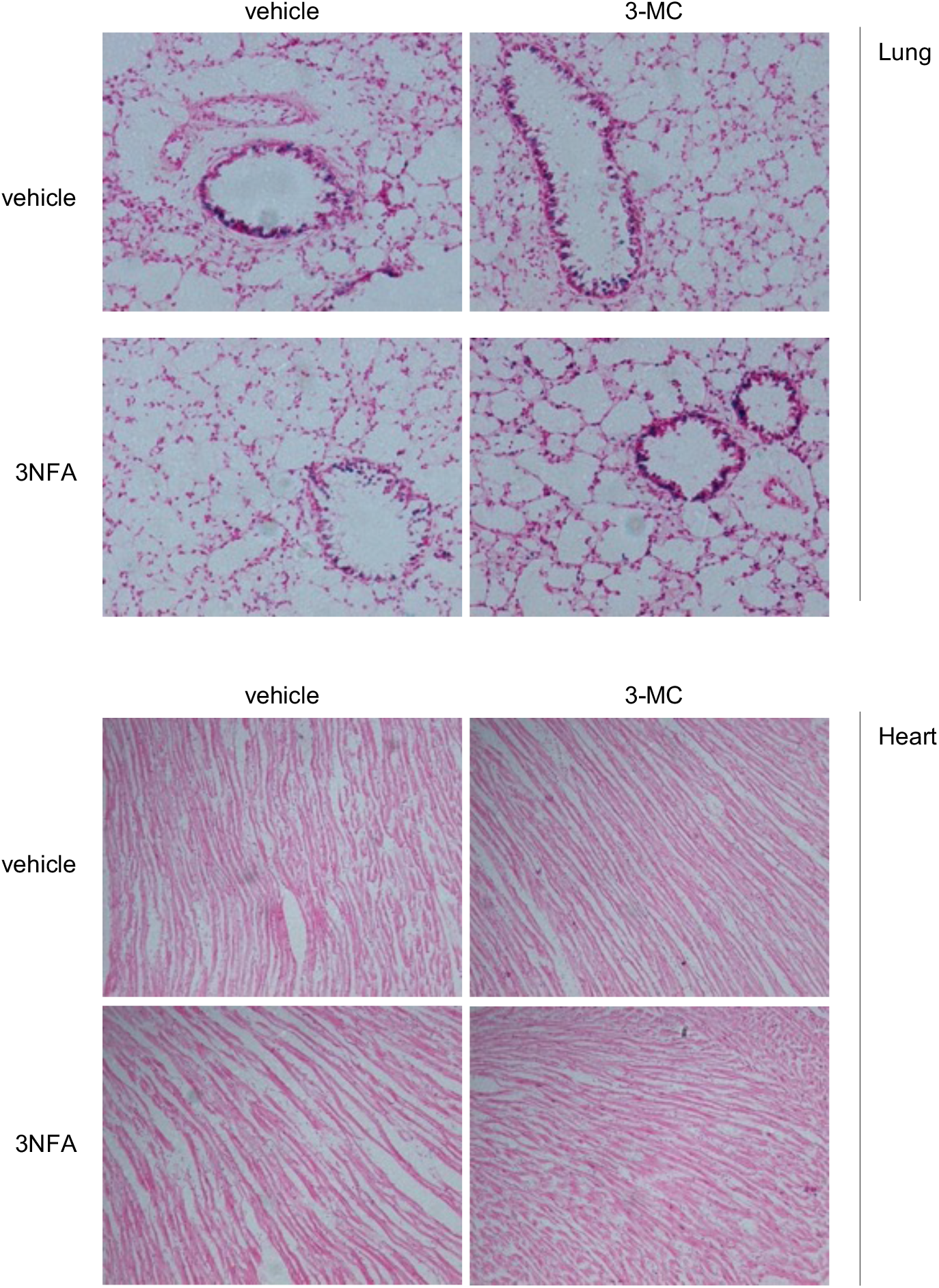
In vivo metabolic activation enhances oxidative stress responses to 3-NFA. Experiment as in Figure 8D, but lung and heart tissues LacZ expression analysis is shown.

